# A maternal-effect genetic incompatibility in *Caenorhabditis elegans*

**DOI:** 10.1101/112524

**Authors:** Eyal Ben-David, Alejandro Burga, Leonid Kruglyak

## Abstract

Selfish genetic elements spread in natural populations and have an important role in genome evolution. We discovered a selfish element causing a genetic incompatibility between strains of the nematode *Caenorhabditis elegans.* The element is made up of *sup-35,* a maternal-effect toxin that kills developing embryos, and *pha-1*, its zygotically expressed antidote. *pha-1* has long been considered essential for pharynx development based on its mutant phenotype, but this phenotype in fact arises from a loss of suppression of *sup-35* toxicity. Inactive copies of the *sup-35/pha-1* element show high sequence divergence from active copies, and phylogenetic reconstruction suggests that they represent ancestral stages in the evolution of the element. Our results suggest that other essential genes identified by genetic screens may turn out to be components of selfish elements.

## Introduction

Selfish genetic elements subvert the laws of Mendelian segregation to promote their own transmission (Dawkins 1976; Doolittle and Sapienza 1980; Orgel and Crick 1980; Werren 2011; Sinkins 2011). In what is perhaps the most extreme scenario, selfish elements can kill individuals that do not inherit them, leading to a genetic incompatibility between carriers and non-carriers (Beeman *et al.* 1992; Werren 1997, 2011;Hurst and Werren 2001; Lorenzen *et al.* 2008). Selfish elements are predicted to spread in natural populations (Hurst and Werren 2001; Werren 2011), and consequently, there is significant interest in using synthetic forms of such elements to drive population replacement of pathogen vectors in the wild (Chen *et al.* 2007; Hammond *et al.* 2015). However, despite the prominent role of genetic incompatibilities in genome evolution and their promise in pathogen control, their underlying genetic mechanisms have been resolved in only a few cases (Werren 2011). Our laboratory previously identified the only known genetic incompatibility in the nematode *Caenorhabditis elegans* (Seidel *et al.* 2008, 2011). The incompatibility is caused by a selfish element composed of two tightly linked genes: *peel-1,* a sperm-delivered toxin, and *zeel-1*, a zygotically expressed antidote. In crosses between isolates that carry the element and ones that do not, the *peel-1* toxin is delivered by the sperm to all progeny, so that only embryos that inherit the element and the *zeel-1* antidote survive. An analogous element, Maternal-effect dominant embryonic arrest (*Medea*) has been previously described in the beetle *Tribolium*; however, the underlying genes remain unknown (Beeman *et al.* 1992; Lorenzen *et al.* 2008).

## Results

### A maternal-effect genetic incompatibility in *C. elegans*

As part of ongoing efforts to study natural genetic variation in *C. elegans*, we introgressed a genetic marker located on the right arm of Chr. V from the standard laboratory strain N2 into the strain DL238 by performing eight rounds of backcrossing and selection. DL238 is a wild strain isolated in the Manuka Natural Reserve, Hawaii, USA, and is one of the most highly divergent *C. elegans* isolates identified to date (Andersen *et al.* 2012). To confirm the success of the introgression, we genotyped the resulting strain at single-nucleotide variants (SNVs) between DL238 and N2 by whole-genome sequencing. As expected, with the exception of a small region on the right arm of Chr. V where the marker is located, most of the genome was homozygous for the DL238 alleles (Fig. 1A). However, to our surprise, we observed sequence reads supporting the N2 allele at many SNVs on Chr. III, including two large regions that were homozygous for the N2 allele despite the eight rounds of backcrossing (Fig. 1A, Fig. S1). This observation suggested that N2 variants located on this chromosome were strongly selected during the backcrossing.

**Figure 1.**
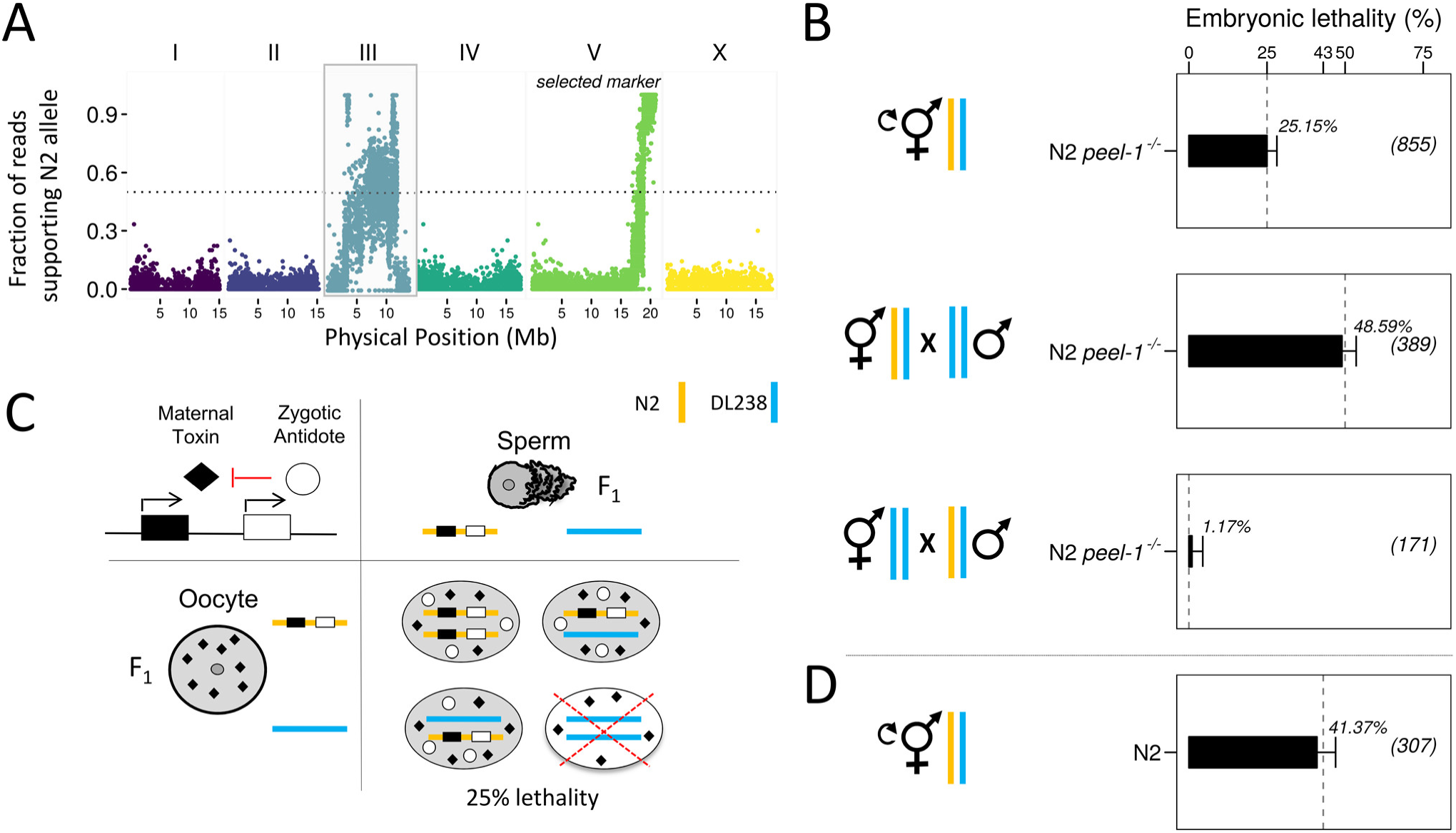
A maternal-effect genetic incompatibility on Chr. III. (A) A marker on Chr. V was introgressed from the reference strain N2 into the DL238 wild isolate. Short-read sequencing of the introgression strain revealed homozygous N2 variants on Chr. III, indicating strong selection in favor of N2 variants during the generation of this strain. **(B)** DL238 males were crossed to hermaphrodites carrying a null allele of the *peel-1/zeel-1* element (*niDfl*) in an otherwise N2 background (N2 *peel-1^-/-^).* F_1_ hermaphrodites were allowed to self-fertilize (top). Alternatively, F_1_ males (middle) or hermaphrodites (bottom) were backcrossed to the DL238 parental strain. Embryonic lethality was scored in the F_2_ progeny as percent of unhatched eggs. Dashed grey lines indicate expected embryonic lethality under the maternal-effect toxin and zygotic antidote model (see also Fig. S2). Sample sizes are shown in parentheses. Error bars indicate 95% binomial confidence intervals, calculated using the Agresti-Coull method (Agresti and Coull 1998)**(C)** Punnett square showing the expected lethality in the F_2_. An interaction between a maternal toxin (black rhombus) and a zygotic antidote (white circle) results in 25% embryonic lethality in the F_2_ and is compatible with the lethality observed in our crosses. (D) Embryonic lethality in the F_2_ progeny of a cross between wild-type N2 hermaphrodites and DL238 males. N2 carries an active copy of *peel-1/zeel-1*, while DL238 carries an inactive copy. Independent segregation of two fully penetrant parental-effect incompatibilities is expected to result in 43.75% embryonic lethality. Orange and blue bars denote N2 and DL238 haplotypes, respectively.

To investigate the nature of the selection, we performed a series of crosses between the N2 and DL238 strains and examined their progeny. To avoid effects of the *peel-1/zeel-1* element, which is present in N2 and absent in DL238, we performed a cross between DL238 males and a near isogenic line (NIL) that lacks the *peel-1/zeel-1* element in an otherwise N2 background (hereafter, N2 *peel-1^-/-^*) (Seidel *et al.* 2011). We observed low baseline embryonic lethality in the F_1_ generation and in the parental strains (0.26% (N *= 381)* for F_1_; 0.99% (*N* = 304) for DL238; 0.4% (*N* = 242) for N2 *peel-1^-/-^*), and we did not observe any obvious abnormal phenotypes in the F1 that could explain the strong selection. However, when we allowed heterozygous F_1_ hermaphrodites from this cross to self-fertilize, we observed 25.15% (N = 855) embryonic lethality among the F_2_ progeny (Fig. 1B). Similar results were obtained for F_1_ hermaphrodites from the reciprocal parental cross (26.1%, *N* = 398). These results suggested the presence of a novel genetic incompatibility between N2 and DL238 that causes embryonic lethality in their F_2_ progeny.

The observed pattern of embryonic lethality (no lethality in the parents nor in the F_1_; 25% lethality in the F_2_) is consistent with an interaction between the genotype of the zygote and a maternal or paternal effect (Fig. 1C) (Seidel *et al.* 2008). We hypothesized that the incompatibility could stem from a cytoplasmically-inherited toxin that kills embryos if they lack a zygotically expressed antidote, analogous to the mechanism of the *peel-1/zeel-1* element (Seidel *et al.* 2008, 2011). To test this model and to discriminate between maternal and paternal effects, we crossed heterozygous F_1_ DL238 x N2 *peel-1^-/-^* males and hermaphrodites with DL238 hermaphrodites or males, respectively (Fig. 1B, Fig. S2). We observed 48.59% (*N =* 389) lethality when F_1_ hermaphrodites were crossed to DL238 males, but only baseline lethality (1.17%; *N* = 171) in the reciprocal cross of F_1_ males to DL238 hermaphrodites. 50% lethality when the F_1_ parent is the mother and no lethality when the F_1_ parent is the father indicates that the incompatibility is caused by maternal-effect toxicity that is rescued by a linked zygotic antidote (Fig. S2). We tested whether the new incompatibility was independent from the paternal-effect *peel-1/zeel-1* element by crossing DL238 and N2 worms and selfing the F_1_ progeny. We observed 41.37% (*N* = 389) embryonic lethality among the F_2_ progeny, consistent with expectation for Mendelian segregation of two independent incompatibilities (43.75%) (Fig. 1D).

### *pha-1* and *sup-35* constitute a selfish element that underlies the incompatibility between DL238 and N2

To identify the genes underlying the maternal-effect incompatibility between N2 and DL238, we sequenced the genome of DL238 using Illumina short reads and aligned those reads to the N2 reference genome. We focused our attention on the two regions on Chr. III that were completely homozygous for the N2 allele in the introgressed strain (Fig. 1A, Fig. S1). Inspection of short read coverage revealed a large ~50 kb region on the right arm of the chromosome with very poor and sparse alignment to the N2 reference (Chr III: 11,086,500 - 11,145,000) (Fig. 2A). This region contains ten genes and two pseudogenes in N2. We noticed that *pha-1,* annotated as an essential gene in the reference genome, appeared to be completely missing in DL238 (Fig. 2A) (Schnabel and Schnabel 1990). *pha-1* was originally identified as an essential gene required for differentiation and morphogenesis of the pharynx, the *C. elegans* feeding organ (Schnabel and Schnabel 1990). But if *pha-1* is essential for embryonic development and missing in DL238, then how are DL238 worms alive? *pha-1* lethality can be fully suppressed by mutations in three other genes: *sup-35, sup-36,* and *sup-37* (Schnabel *et al.* 1991). We found no coding variants in *sup-36* and *sup-37* which reside on chromosomes IV and V, respectively (Schnabel *et al.* 1991) (Fig. S3). However, *sup-35,* which is located 12.5kb upstream of *pha-1*, also appeared to be missing or highly divergent in DL238 (Fig. 2A, Fig. S3).

**Figure 2.**
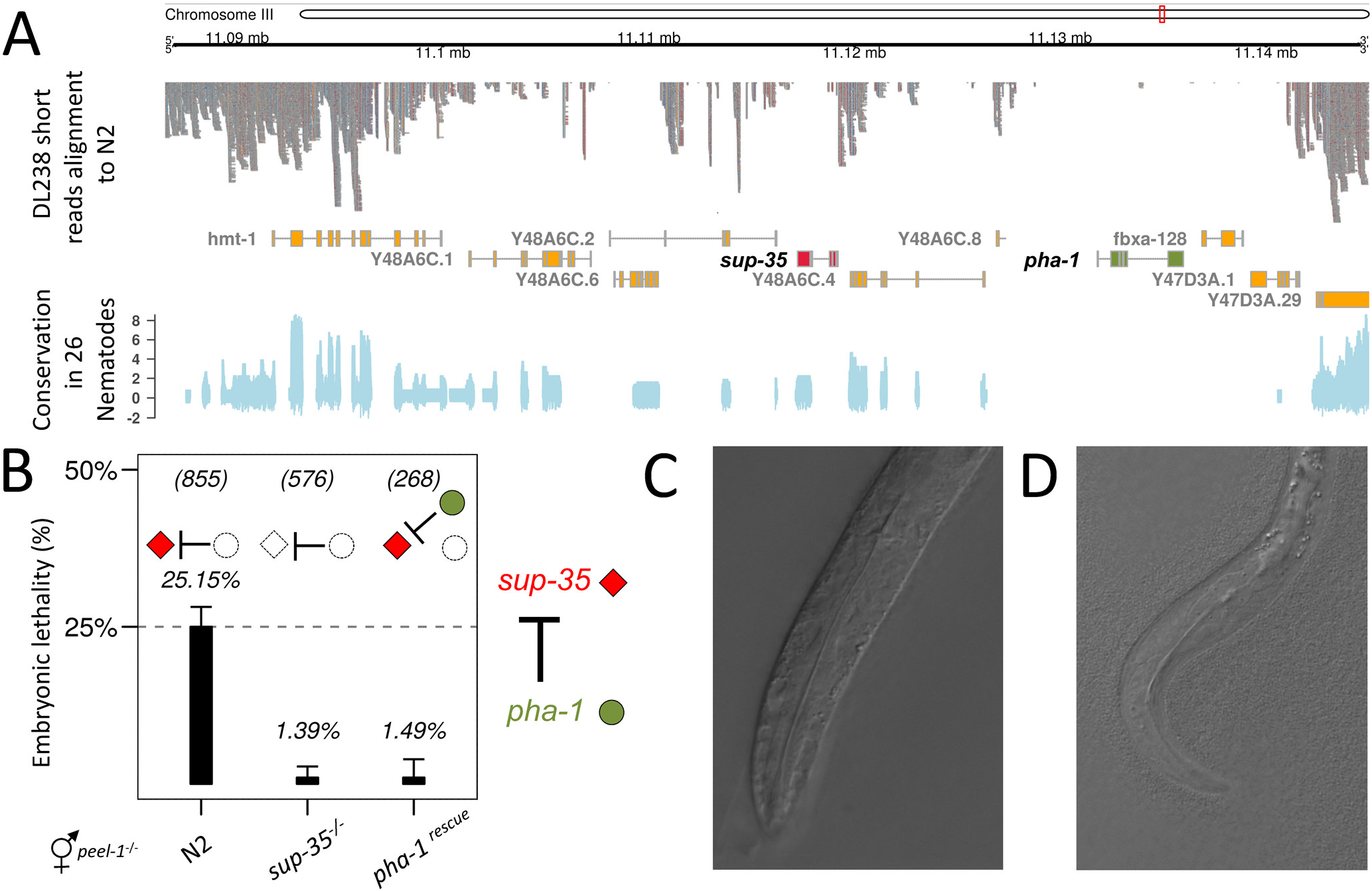
*sup-35* and *pha-1* encode a maternal-effect genetic incompatibility. (A) Alignment of short reads from DL238 to the N2 reference genome (top). A ~50kb region on the right arm of Chr. III selected during the introgression shows sparse alignment throughout with no read support for *pha-1* (green) and weak support for *sup-35* (red). Sequence conservation across 26 nematodes showed no conservation of *pha-1* (bottom). Values are *phyloP* scores retrieved from the UCSC genome browser (Pollard *et al.* 2010) **(B)** In our model, *sup-35* (red rhombus) is a maternally deposited toxin and *pha-1* is a zygotically expressed antidote (green circle). The embryonic lethality in the F_2_ of the cross between DL238 and N2*peel-1^-/-^* (left) was completely rescued when DL238 males were crossed to a strain carrying a *sup-35(e2223)* loss of function allele (center), and also when both parents carried a *pha-1* transgene (right). Error bars indicate 95% binomial confidence intervals, calculated using the Agresti-Coull method (Agresti and Coull 1998)(C) The pharynx of a phenotypically wild-type F_2_ L1 worm from a DL238 x N2 *peel-1^-/-^* cross. **(D)** The pharynx of an F_2_ L1 from the same cross as in (C) showing pharyngeal morphological defects and arrested development.

We hypothesized that *sup-35* and *pha-1* could constitute a selfish element responsible for the observed incompatibility between the N2 and DL238 isolates. In our model, *sup-35* encodes a maternally-deposited toxin that kills embryos unless they express the zygotic antidote, *pha-1* (Fig. 2B). N2 worms carry the *sup-35/pha-1* element, which is missing or inactive in DL238, and F_1_ hermaphrodites deposit the *sup-35* toxin in all their oocytes. 25% of their F_2_ self-progeny do not inherit the element and are killed because they lack the antidote *pha-1.* Consistent with our model, an RNA-sequencing time-course of *C. elegans* embryogenesis showed that *sup-35* transcripts are maternally provided, whereas *pha-1* transcripts are first detected in the embryo at the 100-cell stage (Hashimshony *et al.* 2014). To test our model, we first asked whether *sup-35* was necessary for the F_2_ embryonic lethality in the N2 x DL238 cross. We crossed DL238 males to N2*peel-1^-/-^* hermaphrodites carrying a null *sup-35(e2223)* allele (Fig. 2B). This *sup-35* allele was reported to fully rescue *pha-1* associated embryonic lethality (Schnabel *et al.* 1991). Embryonic lethality in the F_2_ dropped from 25% to baseline in this cross (1.40%, *N = 576*), demonstrating that *sup-35* activity underlies the incompatibility between N2 and DL238 (Fig. 2B). We next tested whether expression of *pha-1*, the zygotic antidote, was sufficient to rescue the embryonic lethality. We introgressed a *pha-1* multicopy transgene into the DL238 and N2 *peel-1^-/-^* strains and repeated the cross. As predicted, expression of *pha-1* was sufficient to reduce embryonic lethality in the F_2_ to baseline (1.49%, *N = 268)* (Fig. 2B). Moreover, we reasoned that if the *sup-35/pha-1* element underlies the maternal incompatibility, arrested embryos from an N2 x DL238 cross should phenocopy *pha-1* mutant embryos. We collected rare L1-arrested F_2_ larvae from an N2 *peel-1^-/-^* x DL238 cross and observed major morphological defects in the pharynx of these individuals, as previously reported for *pha-1* mutants (Schnabel and Schnabel 1990; Polley *et al.* 2014) (Fig. 2C).

Together, these results show that *sup-35* toxicity underlies the incompatibility between N2 and DL238, and that *sup-35* and *pha-1* constitute a selfish element in *C. elegans.* Moreover, our results indicate that *pha-1* is not an organ-specific differentiation gene, as originally proposed (Schnabel and Schnabel 1990), but a zygotically expressed antidote, and *sup-35* is a maternal-effect toxin rather than a suppressor of *pha-1*. This major reinterpretation of the roles of *pha-1* and *sup-35* is strongly supported by multiple lines of evidence from previous studies (Mani and Fay 2009; Fay *et al.* 2012; Kuzmanov *et al.* 2014; Polley *et al.* 2014). First, *sup-35* overexpression phenocopies *pha-1* mutations, showing that *sup-35* is sufficient to cause embryonic lethality (Mani and Fay 2009). Second, all defects associated with *pha-1* mutations are suppressed by mutations in *sup-35* (Mani and Fay 2009; Fay *et al.* 2012; Kuzmanov *et al.* 2014; Polley *et al.* 2014). Third, when N2 hermaphrodites heterozygous for a deletion that includes both *sup-35* and *pha-1 (tDf2/+)* self, the 25% of their progeny that are homozygous for this deletion arrest as embryos with pharyngeal defects (Schnabel *et al.* 1991; Mani and Fay 2009). Importantly, the lethality and pharyngeal defects of those homozygous embryos can be rescued by growing the heterozygous *tDf2/+* mother in *sup-35* RNAi, which depletes *sup-35* transcripts from the germline (Fay *et al.* 2012). These results indicate that maternally deposited *sup-35* is sufficient to kill embryos that lack *pha-1,* which is consistent with the role of *sup-35* as a maternal-effect toxin. Finally, *pha-1* is not found in any other nematode sequenced to date (Fig 2A). This observation is more consistent with its recent evolution as part of a selfish element in *C. elegans* than with its previously postulated role as a key developmental regulator (Schnabel and Schnabel 1990).

### Global variation in the activity of the *sup-35/pha-1* element

We examined the sequences of *sup-35* and *pha-1* in 152 *C. elegans* wild isolates that represented unique isotypes (Cook *et al.* 2016) in the *Caenorhabditis elegans* Natural Diversity Resource (Cook *et al.* 2017). Two isolates, QX1211 (California, USA) and ECA36 (Auckland, New Zealand), harbored a highly mutated copy of *pha-1,* with multiple non-synonymous SNVs as well as frameshifts expected to completely disrupt the protein (Fig. S3 and S4). Both of these isolates also appeared to be missing *sup-35* (Fig. S3). We predicted that these isolates should be incompatible with N2. Because QX1211 and ECA36 carry the same haplotype in the *sup-35 /pha-1* region, we focused on further characterizing QX1211. We crossed QX1211 to N2 *peel-1^-/-^* worms and observed 23.9% (*N* = 355) lethality in the F_2_ progeny, consistent with QX1211 carrying a degenerate copy of *pha-1.* Importantly, the lethality was abolished when we crossed QX1211 with N2 *sup-35(e2223);peel-1^-/-^* (0%, *N* = 290). Furthermore, we observed background levels of embryonic lethality (1.0%, *N* = 294) in the F_2_ progeny of a DL238 x QX1211 cross, as expected, since both strains lack functional *sup-35.* Our analysis also revealed that the highly divergent Hawaiian isolate CB4856 carried a functional *sup-35*/*pha-1* element, which explains why previous studies did not detect this incompatibility (Seidel *et al.* 2008). Consistent with this observation, crossing the CB4856 and DL238 isolates led to the expected embryonic lethality in the F_2_ (22.3%, *N = 349*).

We looked for additional variation in *sup-*35, *sup-*36, and *sup-*37 across the 152 isolates, which could potentially affect the activity of the *sup-35/pha-1* element (Fig. S5-7). We found eight non-synonymous variants (three in *sup-*35, three in *sup-*36 and two in *sup-37*) and one premature stop codon in *sup-35* that removed 47% of the protein. We also identified potential deletions by visually inspecting read alignments in each of the 152 isolates. While *sup-36* and *sup-*37 had consistent coverage in all isolates, we identified two structural variants in *sup-*35: a 530 bp deletion in the third intron, and a large 12.1 kb deletion that removed part of the last exon and the 3’ UTR and fused *sup-35* to *Y48A6C.1*, a pseudogene that has partial homology with *sup-35,* creating a chimeric transcript (Fig. S8 and S9). We tested strains carrying each of these variants for a maternal-effect incompatibility with DL238 (Table S2, Fig. S10), and found that the incompatibility was completely abolished in strains carrying the chimeric *sup-35/Y48A6C.1* gene and in the strain carrying the premature stop codon in *sup-35*, indicating that these variants disrupt *sup-35* function. Thus loss of *sup-35* activity has occurred independently at least twice in carriers of N2-like alleles the element.

### DL238 and QX1211 carry an ancestral *sup-35/pha-1* haplotype

The alignment of DL238 and QX1211 short reads to the N2 reference genome was very sparse throughout the *sup-35/pha-1* region and at nearby genes, with some genes aligning only in exons and others not aligning at all (Fig. 2A). Moreover, several attempts to define the boundaries of the *pha-1* deletion in DL238 using diverse combinations of PCR primer pairs were unsuccessful. This suggested that the DL238 and QX1211 haplotypes were highly divergent from the N2 reference, and that major genomic rearrangements may have occurred. To resolve the genomic structure of the *sup-35/pha-1* element in these isolates, we *de novo* assembled the genomes of DL238 and QX1211 using a combination of our and previously published Illumina short reads (van Schendel *et al.* 2015; Cook *et al.* 2017), followed by targeted Sanger sequencing to resolve repetitive regions and confirm scaffolds. The *de novo* assemblies confirmed that *pha-1* is absent from DL238 and is highly pseudogenized in QX1211, and that *sup-35* is pseudogenized in both (Fig. 3, Fig. S11). DL238 and QX1211 share a very similar haplotype, with the exception of a large deletion in DL238 that encompasses *pha-1, fbxa-128* and several exons of *Y47D3A.1* (Fig. S11). We also identified other large structural variants in both DL238 and QX1211 at the *sup-35/pha-1* locus. First, relative to the N2 reference genome, nearly 20kb of sequence is missing completely from both isolates (Fig. S11). Second, the region spanning the pseudogenized *sup-35* and *Y48A6C.4* is inverted relative to the N2 reference (Fig. 3, Fig. S11). This inversion was confirmed using single molecule Oxford Nanopore long-read sequencing (Fig. S12). As a consequence of the inversion, the pseudogenized *sup-35* and *pha-1* are located next to each other in QX1211, rather than flanking *Y48A6C.4* as in the N2 reference genome (Fig. 3, Fig. S11).

**Figure 3.**
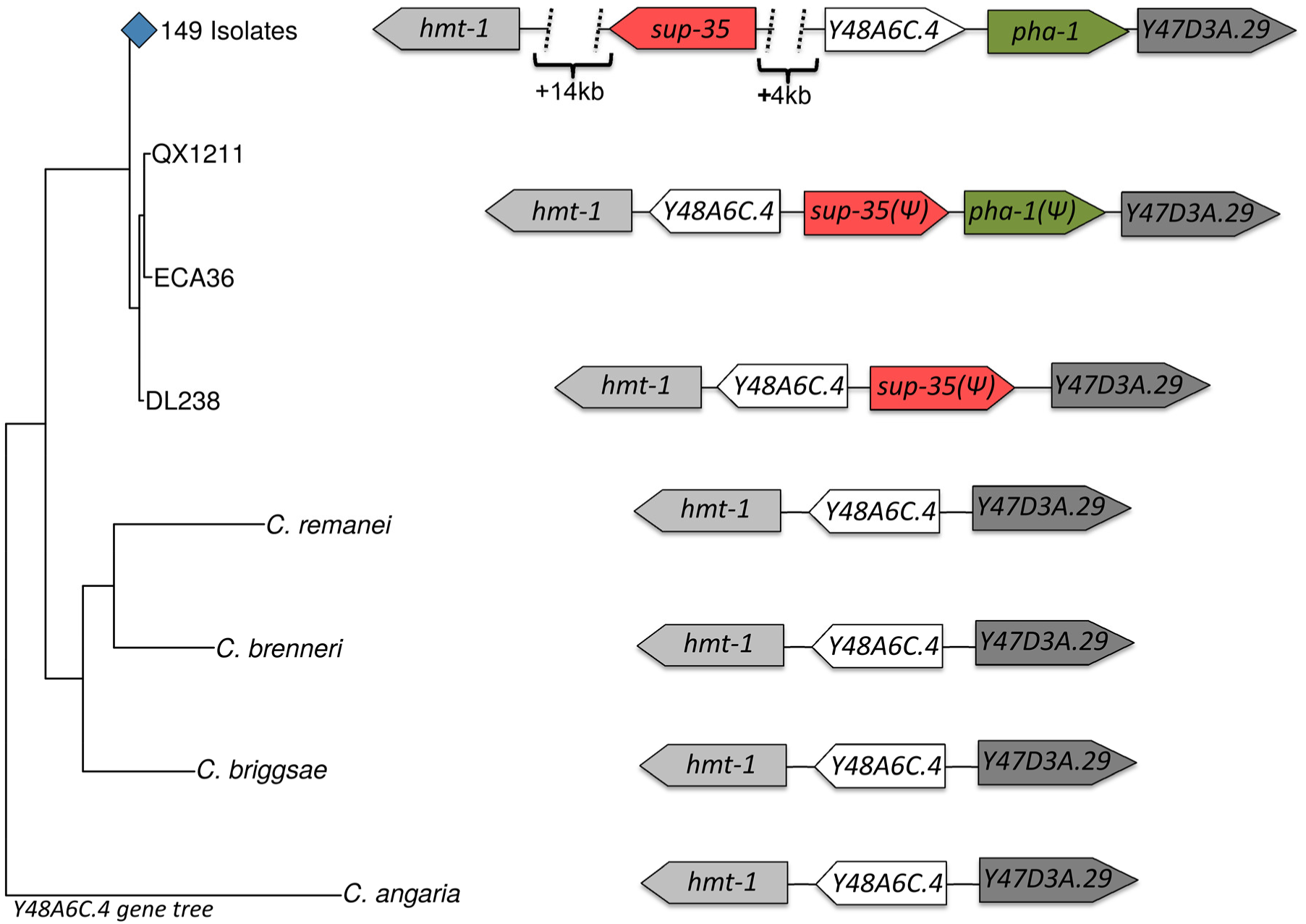
The *sup-35/pha-1* N2 haplotype is derived and is marked by an inversion. (Left) A gene tree built from the coding region of *Y48A6C.4* in 152 *C. elegans* isolates and four other *Caenorhabditis* species. DL238, QX1211 and ECA36 cluster together in a separate branch from all other *C. elegans* isolates. (Right) The synteny in the region containing the *sup-35/pha-1* element, as well as three highly conserved genes in the close vicinity (*hmt-1, Y48A6C.4,* and *Y47D3A.29*) is schematically represented. ***(ψ)*** denotes alleles that are pseudogenized. The genes *sup-35* (red) and *Y48A6C.4* (white) are inverted in DL238, QX1211, and ECA36 relative to the other 149 *C. elegans* isolates. The gene order and orientation of *hmt-1, Y48A6C.4,* and *Y47D3A.29* in other *Caenorhabditis* species suggests that the inverted haplotype is the ancestral, and that the haplotype found in 149 isolates is the derived one.

To gain further insights into evolution ofthe *sup-35/pha-1* element, we aligned the N2, DL238 and QX1211 haplotypes to the homologous regions of diverse *Caenorhabditis* species, using the highly conserved genes (*hmt-1, Y48A6C.4,* and *Y47D3A.29*) that delineated the region (Fig. 3, Fig. S11). Unexpectedly, our analysis revealed that the order and orientation of these three genes in the other *Caenorhabditis* species matched that in DL238 and QX1211 rather than the order and orientation in N2. This observation suggests that the sup-35/pha-1 haplotype in DL238 and QX1211 derives from an early stage in the evolution of the selfish element, which was followed by a major inversion that now defines the N2 haplotype, and subsequent degeneration of the element in DL238 and QX1211. In further support of this model, a gene tree built using the coding region of *Y48A6C.4* from all the *C. elegans* isolates and the other *Caenorhabditis* species showed that DL238, QX1211 and ECA36 cluster in a separate branch from all other *C. elegans* isolates (Fig. 3).

## Discussion

We discovered a genetic incompatibility in *C. elegans* that is caused by an interaction between a maternally deposited toxin, *sup-35*, and a zygotically expressed antidote, *pha-1.* The antidote, *pha-1,* was originally thought to be a developmental gene, in large part due to the specific pharyngeal defects observed in mutants (Schnabel and Schnabel 1990; Granato *et al.* 1994; Okkema *et al.* 1997; Fay *et al.* 2004; Mani and Fay 2009). However, the precise role of *pha-1* in embryonic development remained elusive and controversial (Mango 2009; Fay *et al.* 2012; Kuzmanov *et al.* 2014). Our results indicate that *pha-1* pharyngeal defects are a direct consequence of *sup-35* toxicity, and that *sup-35* and *pha-1* act as a selfish element, instead of being integral components of *C. elegans* embryonic development as originally suggested.

One important insight emanating from previous work in light of our results is that the *sup-35/pha-1* element exerts its toxicity by recruiting genes that are directly involved in *C. elegans* development (Schnabel *et al.* 1991; Fay *et al.* 2004, 2012;Mani and Fay 2009; Polley *et al.* 2014). The other two known suppressors of *pha-1* lethality, *sup-36* and *sup-37,* are essential for *sup-35* toxicity and are conserved in other nematodes (Fay *et al.* 2012; Polley *et al.* 2014). Interestingly, *sup-37* is required for normal pharyngeal pumping and promotes ovulation in the somatic gonad independently of *pha-1* function (Fay *et al.* 2012). Null *sup-37* mutants are inviable and undergo early larval arrest. However, a single missense and viable mutation in *sup-37* is sufficient to abolish *sup-35* toxicity (Fay *et al.* 2012; Polley *et al.* 2014). Together with the finding that SUP-37 physically interacts with SUP-35 (Polley *et al.* 2014), this suggests that the *sup-35/pha-1* selfish element is hijacking a developmental pathway to kill those embryos that do not inherit it. The specificity in the activity and expression of *sup-36* and *sup-37* may explain the pharyngeal phenotypes of *pha-1* mutants. We hypothesize that PHA-1 could act as an antidote by directly inhibiting the interaction between SUP-35 and SUP-37. The transcription factor *lin-35/Rb* and the E2 ubiquitin conjugation enzyme *ubc-18* downregulate *sup-35* (Mani and Fay 2009). An attractive possibility is that this regulation evolved as an additional mechanism to cope with *sup-35* toxicity, as part of an arms race between the selfish element and its host. Future studies may further resolve the mechanism of *sup-35* toxicity and its regulation.

One of the most intriguing aspects of toxin-antidote systems is their origin. The study of the *pha-1/sup-35* element provides some clues. *pha-1* has no homology to any other gene outside *C. elegans.* On the other hand, *sup-35* is a homolog of another *C. elegans* gene, *rmd-2,* which is conserved in other nematodes (Mani and Fay 2009). A phylogenetic analysis shows that *sup-35* is more closely related to *C. elegans rmd-2* than to *rmd-2* genes from other species, and is likely a paralog of *rmd-2* (Fig. S13 and S14). These results suggest that the origin of the *sup-35/pha-1* element involved the duplication of a pre-existing gene *(rmd-2)* and the recruitment of a novel gene of unknown origin in the lineage leading to *C. elegans.*

Among 152 *C. elegans* wild isolates examined, only DL238, QX1211, and ECA36 do not carry the derived inversion in the *sup-35/pha-1* element, and in all three of them, the selfish element is highly pseudogenized. Similar inversions have been described in the *Drosophila* segregation distorter locus and in the mouse *t-haplotypes* (Shin *et al.* 1983; Hurst and Werren 2001; Larracuente and Presgraves 2012), and are thought to stabilize two-component driver systems by preventing recombination from decoupling the components (Hurst and Werren 2001). Has the inversion facilitated the spread of *sup-35/pha-1* through the *C. elegans* population to all but few isolates? Ongoing efforts to identify more divergent isolates, as well as nematode species that are more closely related to *C. elegans,* may fill in the gaps in our understanding of the evolution of this element.

Lastly, our work highlights the importance of studying natural genetic variation for understanding gene function. Despite the indisputable value of a common reference strain, it has proved extremely difficult in the context of the N2 background alone to either confirm or rule out *pha-1* as an essential component of *C. elegans* embryonic development. The study of other wild isolates has made possible our characterization of *sup-35/pha-1* as a selfish element. Our results show that some essential genes may, in fact, turn out to be antidotes to unknown toxins. Selfish elements conferring genetic incompatibilities may be more common than previously thought, and some of them may be hiding in plain sight.

## Acknowledgments

We thank members of the Kruglyak lab for their comments. Funding was provided by the Howard Hughes Medical Institute and NIH grant R01 HG004321 (L.K.). E.B. is supported by a Gruss-Lipper postdoctoral fellowship from the EGL foundation. A.B. is supported by the Jane Coffin Childs Memorial Fund for Medical Research. E.B., AB. and L.K. wrote the manuscript. All authors discussed and agreed on the final version of the manuscript. The authors declare no competing financial interests.

## Methods

### *C. elegans* strains and mutant alleles

Strains were grown using standard culturing techniques, with the exception that a modified nematode growth medium (NGM) containing 1% agar and 0.7% agarose was used to prevent burrowing of wild isolates (Brenner 1974; Andersen *et al.* 2015). All experiments were carried out at 20^o^C. All the strains used and generated in this study are listed in Table S2. Some of the strains were provided by the CGC, which is funded by the NIH Office of Research Infrastructure Programs (P40 OD010440). The construction of strains carrying the *peel-1/zeel-1* allele from CB4856 (*niDf9*) was performed by backcross following a PCR product specific to the N2 allele amplified using the following primers: FW GCAGAGGAGGCAAAGGTGACTA; RV AGCACGTGT AGGCAGAAGTCAT.

### Introgression of a Chr. V genetic marker into DL238

We introgressed the *fog-2* (*q71*) allele from the N2 background into DL238 by performing eight consecutive rounds of backcross and selection for the *feminization of the germline (fog)* phenotype (Schedl and Kimble 1988). Since the *peel-1/zeel-1* element is active in N2 but not in DL238, we performed the backcross using DL238 males and feminized hermaphrodites. The use of DL238 males avoided the fixation of the *peel-1/zeel-1* element on Chr. I because DL238 worms do not carry the PEEL-1 toxin in their sperm. Consequently, the fixation of the N2 haplotype on Chr. III in the introgression strain could not be caused a paternal-effect toxin. The introgressed marker, *fog-2(q71)*, is required for spermatogenesis in hermaphrodites but not in males, and the resulting strain must reproduce by outcrossing. The observed pattern of linkage decay around the N2 homozygous region on Chr. III (Fig. S1) is a consequence of selection acting in an obligate outcrossing population.

### Short read sequencing

We extracted genomic DNA (gDNA) using the DNeasy Blood & Tissue kit (Qiagen). We prepared Illumina sequencing libraries using the Nextera protocol (Illumina). We followed the standard protocol with the following exception: we performed agarose size-selection of the Nextera libraries, extracting a ~500 bp band. Libraries were quantified using the Qubit HS kit and sequenced using 300bp paired-end V3 kits on a Miseq desktop sequencer (Illumina) at 12 pM. In total, we generated 29,998,002 reads for DL238 and 14,428,754 reads for QX2323.

### Variant calling in DL238

Automated preprocessing, alignment and variant calling were done using *bcbio-nextgen* (*ver. 0.9.9*) (*https://bcbio-nextgen.readthedocs.io/*). Short reads from DL238 were aligned to the WBcel235 build of the reference N2 genome. Variant calling was performed with four different software packages: *GATK HaplotypeCaller* (McKenna *et al.* 2010), *Platypus* (Rimmer *et al.* 2014), *Freebayes* (Garrison and Marth 2012) and *Varscan* (Koboldt *et al.* 2012). Variants identified by fewer than three of the callers were filtered out, resulting in 239,131 SNVs between DL238 and N2.

### Genotyping of the DL238 isolate and the DL238 Chr. V introgression strain

Analyses were performed using the R Project for Statistical Genetics (https://www.r-project.org/). Plotting was done using the R package *ggplot2* (Wickham 2009) and genomic data was plotted using the R package *gviz* (Hahne and Ivanek 2016). We sought to accurately assess the success of the introgression genome-wide, rather than exhaustively identify variants. Therefore, we implemented a highly stringent filtering pipeline to extract only the highest confidence SNVs from our initial list of 239,131. First, we used all variants (SNVs and Indels) to generate an alternative DL238 reference build. We extracted a 50bp region around each SNV from the N2 reference genome and aligned it to the DL238 alternative reference using *bowtie (ver. 1)* (Langmead *et al.* 2009), allowing up to 5 mismatches. This was implemented using the R package *liftoveR* we have developed (https://github.com/eyalbenda/liftoveR), and allowed us to eliminate SNVs that were proximal to structural variants (which wouldn’t align with *bowtie).* That reduced our set to 141,073 SNVs. We aligned both the QX2323 and the DL238 sequencing reads using *burrows wheeler aligner (bwa)* (Li and Durbin 2010) to the N2 and the DL238 references, and removed PCR duplicates using *Picard.* We then counted the number of reads supporting each allele of each of the 141,073 SNVs identified in DL238 using *bam-readcount (https://github.com/genome/bam-readcount).* To reduce the number of spurious SNVs, we restricted our analysis to SNVs for which all DL238 reads supported the DL238 allele when aligning to DL238, and no reads supported the N2 allele when aligning to N2. This reduced the number of SNVs to 53,186. Finally, to accurately assess the dosage of each allele, SNVs with less than 20x coverage were filtered out. 44,200 SNVs passed filtering. The fraction of reads with the N2 allele at these SNVs was plotted without smoothing.

### Mating and embryonic lethality scoring in the crosses between DL238 and N2 *peel-1^-/-^*

Males and L4 hermaphrodites were picked onto an NGM plate seeded with a small drop of OP50 bacteria and left to mate. We used a high male to hermaphrodite ratio (at least 5:1), and inspected the F_1_ progeny to ensure both sexes were equally represented. To score embryonic lethality in the progeny of selfing F_1_ hermaphrodites, F_1_ L4 hermaphrodites were transferred to a new plate. Next morning, gravid hermaphrodites were allowed to lay eggs for 4-8 hours. Eggs laid during this time window were collected and manually transferred to a new unseeded plate, counted and scored 18-20 hours later. Lethality was defined as the percentage of unhatched eggs. Embryonic lethality in the progeny of mating hermaphrodites was scored similarly, but L4 hermaphrodites were transferred together with males to a new plate, and their eggs were laid and collected in the presence of these males to guarantee continuous mating.

### Mapping of the genetic incompatibility

To identify the locus underlying the strong selection in favor of the N2 haplotype on Chr. III, we first identified regions homozygous for the N2 alleles in the QX2323 introgression strain using our curated SNV list. We identified two regions: Chr III. 3,302,436–3,607,723 and Chr. III. 10,964,134–11,254,095 (Fig. S1). Of these two regions, only the one located toward the right arm of Chr. III showed a pattern of decaying linkage, suggesting that this region was more likely to harbor the causal locus. The homozygous N2 region on the left arm of Chr. III could be the result of an unknown structural variant in DL238 unrelated to the incompatibility. Visual inspection of DL238 short read alignments in the homozygous region revealed that *pha-1* and *sup-35* were very likely missing in DL238. To test whether *sup-35* was necessary for the genetic incompatibility, we used the GE346 strain carrying the *sup-35(e2223)* allele that was shown to suppress *pha-1* mutations (Schnabel *et al.* 1991). The strain also carried in close linkage a temperature sensitive *(ts) pha-1(e2123)* allele, as well as a recessive *dpy-18(e499)* mutation that caused a dumpy *(dpy)* phenotype. All crosses were performed at 20°, the permissive temperature of the *pha-1(e2123)* allele. F_1_ cross progeny from *dpy* hermaphrodites were confirmed by lack of the *dpy* phenotype. Since GE346 carries the active N2 *peel-1/zeel-1* allele, we crossed GE346 into the N2 *peel-1^-/-^* strain, selecting for *dpy* and the absence of *peel-1.* Hermaphrodites of the resulting strain (QX2329) were crossed to DL238 males. To test whether transgenic expression of *pha-1* was sufficient to rescue the embryonic lethality, we used the OH10050 strain carrying the integrated extrachromosomal array *otIs317* [*mgl-1::mCherry* + *pha-1* (+)]. We introgressed the transgene from OH10050 into DL238 by performing eight rounds of backcross and selection for *mCherry* fluorescence. We also introgressed the transgene into the N2 *peel-1^-/-^* strain, and crossed the two strains. This guaranteed expression of the transgene in all F_2_ progeny.

### Microscopy

We selected phenotypically WT and rare L1 arrested embryos from the F_2_ of an N2 *peel-1^-/-^* x DL238 cross. Larvae were transferred to a 3% agarose pad and visualized under bright field using a Nikon Eclipse 90i microscope equipped with a Photometrics CoolSNAP HQ2 CCD camera.

### Screening for variants in *sup-35, sup-36* and *sup-37* across wild isolates

Raw Illumina short reads sequencing data from 152 isolates (Cook *et al.* 2017) was acquired from the Sequence Read Archive (SRA) (Bioproject PRJNA318647). Alignment and variant calling were done using *bcbio-nextgen* as described above for DL238 to identify coding variants in *sup-35., sup-36* or *sup-37.* To identify structural variation, we visualized the alignment in each of these three genes across the isolates. finding two structural variants in *sup-35* (Fig. S5.S8 and S9). The variants were visualized using mutation “lollipop” plots generated using the *trackViewer* Bioconductor R package. Multiple sequence alignments were visualized using *Geneious ver. 10 (restricted free version) (https://www.geneious.com/).*

### Screening for *sup-35/pha-1* activity in isolates carrying variants in *sup-35, sup-36* or *sup-37*

To facilitate screening wild isolates for variants that abolish the incompatibility with DL238, we used a *fog-2(q71)* introgression strain in the DL238 background (QX2327) that was homozygous for the DL238 *sup-35/pha-1* allele. This strain was generated by backcrossing the QX2323 strain into DL238 for two additional generations using QX2323 males and DL238 hermaphrodites, thus avoiding the maternal incompatibility. The QX2327 strain was genotyped to ensure it was homozygous for the DL238 *sup-35/pha-1* allele. This strain is obligate outcrossing, and we crossed QX2327 hermaphrodites to males from each of the wild isolates, thereby ensuring that the progeny in the next generation resulted exclusively from crossing and not selfing. We noted that the baseline lethality in this strain was markedly higher than wild-type (4.7%, *N* = 252). However, when we backcrossed QX2327 to DL238, and allowed the F_1_ to self, the lethality was reduced back to background levels (0.5%, *N* = 362). These results explain why higher lethality (>50%) is observed in crosses in which F_1_ males are backcrossed to QX2327 hermaphrodites, while the expected lethality is observed in the selfing F_1_ and in the reciprocal backcrosses that use F_1_ hermaphrodites (Table S1).

In a screen for functional mutations in *sup-35*, we expect null *sup-35* alleles to reduce the lethality from 25% to baseline levels in the F_2_. In contrast, even null variants in *sup-36* or *sup-37,* which are unlinked to *sup-35/pha-1*, are predicted to only reduce the embryonic lethality from 25% to 18.75% if they are recessive. This is due to the fact that of the F_2_ progeny that do not inherit *pha-1* (25%), 75% will inherit at least one functioning copy of the *sup-36* (or *sup-37*) allele. Furthermore, in a backcross of F_1_ males to QX2327 hermaphrodites, all of the F_2_ progeny inherit at least one functioning copy of *sup-36* (or *sup-37*) from the QX2327 parent, and the lethality isn’t expected to be reduced at all. For these reasons, and given the sample sizes in our screening, we cannot rule out the presence of hypomorphic variants weakly affecting *sup-35*, or even strongly affecting *sup-36* and *sup-37*, in some of the wild isolates tested.

### Validation of chimeric fusion

To validate the large deletion leading to the fusion of *sup-35* and the *Y48A6C.1* pseudogene, we designed primers that flanked the deletion: FW-del: GATCACGTGAGACAGGAAAAG and RV-del: CCCTTCAAAAGCACACCAAC. This primer pair amplified the expected 1000 bp band in the wild isolate ED3012, which carries the deletion, but not in the reference N2 (Fig. S8C). As a positive control, we amplified the *pha-1* locus using primers FW-pha-1: CCGTTTTCATCACGTTGCTCGA and RV-pha-1: TGTCGCGCACTACTGAATCAGA. To confirm whether the chimeric fusion *sup-35/Y48A6C.1* is expressed, we performed reverse transcription (RT) PCR (Fig. S8D). Total RNA was isolated from mixed stage N2 and ED3012 populations using the RNeasy kit (Qiagen), and cDNA was prepared using the SuperScript III Reverse Transcriptase kit (Thermo Fisher Scientific). We used primers FW1: TTTTTCGCTTTCCAAACTGG, RV1: GCGAGCAACTCTTTCTCGAT, RV2: ATTTTGAGAGCAAGCCGAAA. The FW1-RV1 primer pair amplifies exclusively the spliced cDNA of wild type *sup-35,* and the FW1-RV2 primer pair amplifies exclusively the spliced cDNA of the *sup-35/Y48A6C.1* chimeric fusion. As a positive control, we amplified the *pha-1* transcript using the primers FW-pha-1.2: CGGACCAGTTCAAAATGACA and RV-pha-1.2: TTTTCTGCTGGGAGATTTGC. The primers span multiple exons, allowing us to distinguish the spliced cDNA PCR product.

### *De novo* genome assembly

We *de novo* assembled the DL238 and QX1211 genomes by combining Illumina short reads generated by two previous studies available at the Sequence Read Archive (Bioproject PRJNA318647 and PRJNA260487) (van Schendel *et al.* 2015; Cook *et al.* 2016). Average insert size in those libraries was 200 bp. In total, we used 113,580,009 100 bp paired-end reads from DL238 (~227X coverage) and 81,268,657 100 bp paired-end reads from QX1211 (~163X coverage). We assembled the genomes using *SOAPdenovo2* (Luo *et al.* 2012) with a K-mer size of 23. We then used *blastn* (Altschul *et al.* 1990) to search for scaffolds that had homology to *sup-* 35, *pha-1,* and genes in their vicinity. The genomic region that contained the *sup-35/pha-1* element and neighboring genes (*hmt-1* and *Y47D3A.29*) was not recovered in a single scaffold because our assemblies lacked mate-pair information. Moreover, the region of interest contained several repetitive elements. To circumvent this limitation and generate a single scaffold, we generated a *de novo* assembly using an additional software package, *DiscovarDenovo (v52488),* with default parameters (Weisenfeld *et al.* 2014). We then used targeted PCR amplification followed by Sanger sequencing to confirm the new scaffolds, and to close remaining gaps. Finally, DL238 and QX1211 Illumina short reads were re-aligned to the assembled haplotypes and visually inspected to discard errors introduced by the assemblers.

### Nanopore single molecule sequencing

We extracted fresh DL238 gDNA using the DNeasy Blood & Tissue kit (Qiagen) and prepared the sequencing library the same day of extraction following the 1D Genomic DNA by ligation (SQK-LSK108) protocol (Oxford Nanopore Technologies) starting from 1.5 μg of gDNA and avoiding DNA fragmentation. Sequencing was performed on a MinION Mk1b using the R9.4 chemistry, and reads were collected for 48 hours. In total, we recovered 194,578 reads that passed quality control (*MinKNOW* software). Those reads were converted to fastq format using *poretools* (Loman and Quinlan 2014). To determine whether the long reads supported our *de novo* assembled DL238 haplotype, we aligned the reads to a modified WBcel235 N2 reference genome that included the DL238 haplotype as an additional scaffold. This allowed us to directly compare the support for either the N2 reference or our *de novo* assembled haplotype. We aligned the nanopore reads using *bwa* with the *ont2d* flag to allow for the high error rate in the reads. The alignment rate was 94.6%. Average read length was 3,962 bp, and 16,110 reads (11%) were over 10 kb in length.

### Multiple sequence alignment

To study the genomic architecture and evolution of the *sup-35/pha-1* element, we recovered the sequence at the *sup-35/pha-1* locus in additional *Caenorhabditis* species. Although the genes themselves are not conserved in any other sequenced nematode species, three genes in close vicinity, *hmt-1*, *Y48A6C.4,* and *Y47D3A.29,* are highly conserved. *hmt-1* is a predicted transmembrane ABC transporter, and *Y47D3A.29* encodes the catalytic subunit of DNA polymerase alpha. *hmt-1* and *Y47D3A.29* flank the *sup-35/pha-1* element in *C. elegans. Y48A6C.4,* a predicted ortholog of *S. cerevisiae* IPI1, is located between *hmt-1* and *Y47D3A.29* in all sequenced *Caenorhabditis.* We took advantage of the high evolutionary conservation of these three genes to recover the sequence of the region in four nematode species (C. *briggsae “cb4”, C. brenneri* “6.0.1b”, *C. remanei* “ASM14951v1”, and *C. angaria* “ps1010rel4”). Those sequences, together with three *C. elegans* haplotypes (N2, DL238 and QX1211), were than aligned using *progressiveMauve* (Darling *et al.* 2010).

### Phylogenetic tree

The coding sequence of *Y48A6C.4* was recovered in the different *C. elegans* isolates and other *Caenorhabditis* species. To determine the sequence of the gene in the possible presence of SNVs or indels, we first generated an alternative reference sequence for each of the isolates based on the SNVs and indels identified in our variant calling analysis. We then assembled the cDNA sequences of *Y48A6C.4* in each isolate. To that end, we first aligned the sequence around the start and end positions of each exon to the alternative reference using *blast* (Altschul *et al.* 1990). Following the alignment, the positions were extracted from the reference and concatenated. This procedure allowed us to ensure the correct gene sequence in the presence of SNVs or indels. Alignment of positions from one genome to another using *blast* is implemented in our *liftoveR* R package (https://github.com/eyalbenda/liftoveR). The cDNA sequences of *Y48AC6.4* orthologs from other *Caenorhabditis* were acquired from the *Wormbase* release WS256. To create a gene tree, the cDNA sequences were *in-silico* translated, and the protein sequences were aligned using *MAFFT* (Katoh and Standley 2013). Codon alignment was then generated from the cDNA sequences and the aligned protein sequences using *pal2nal* (Suyama *et al.* 2006). Finally, the codon-aligned cDNA sequences were used as input for maximum likelihood phylogenetic reconstruction using *MrBayes* (4 chains of 1,000,000 cycles) (Ronquist and Huelsenbeck 2003). The general time reversible (GTR) substitution model with gamma distributed rate variation was used as it was found to be the best fit (by minimizing the BIC criterion) using *jModelTest2* (Darriba *et al.* 2012). The resulting tree was visualized using the R package *ggtree* (Yu *et al.* 2017).

## Supplementary Figures

**Figure S1.**
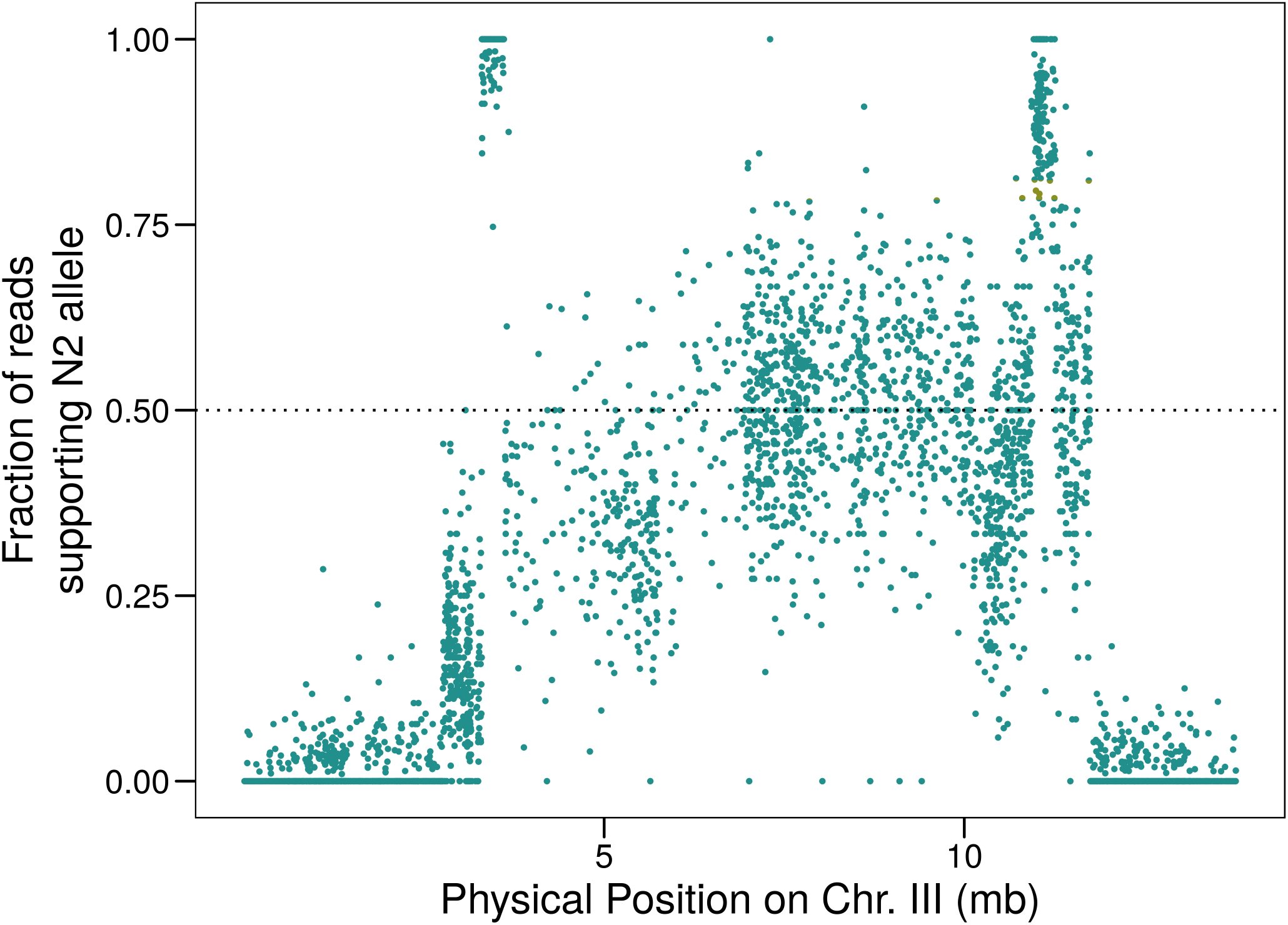
Selection for N2 alleles on Chr. III when introgressing a genetic marker on Chr. V from N2 into DL238. Each point is an SNV that was identified in DL238. A strict pipeline was used to reduce the incidence of spurious SNVs (see Materials and Methods). We identified two regions homozygous for N2 alleles: Chr III. 3,302,436–3,607,723 and Chr. III. 10,964,134–11,254,095. The region closer to the right arm shows linkage decay, suggesting it harbors the locus under selection.

**Figure S2.**
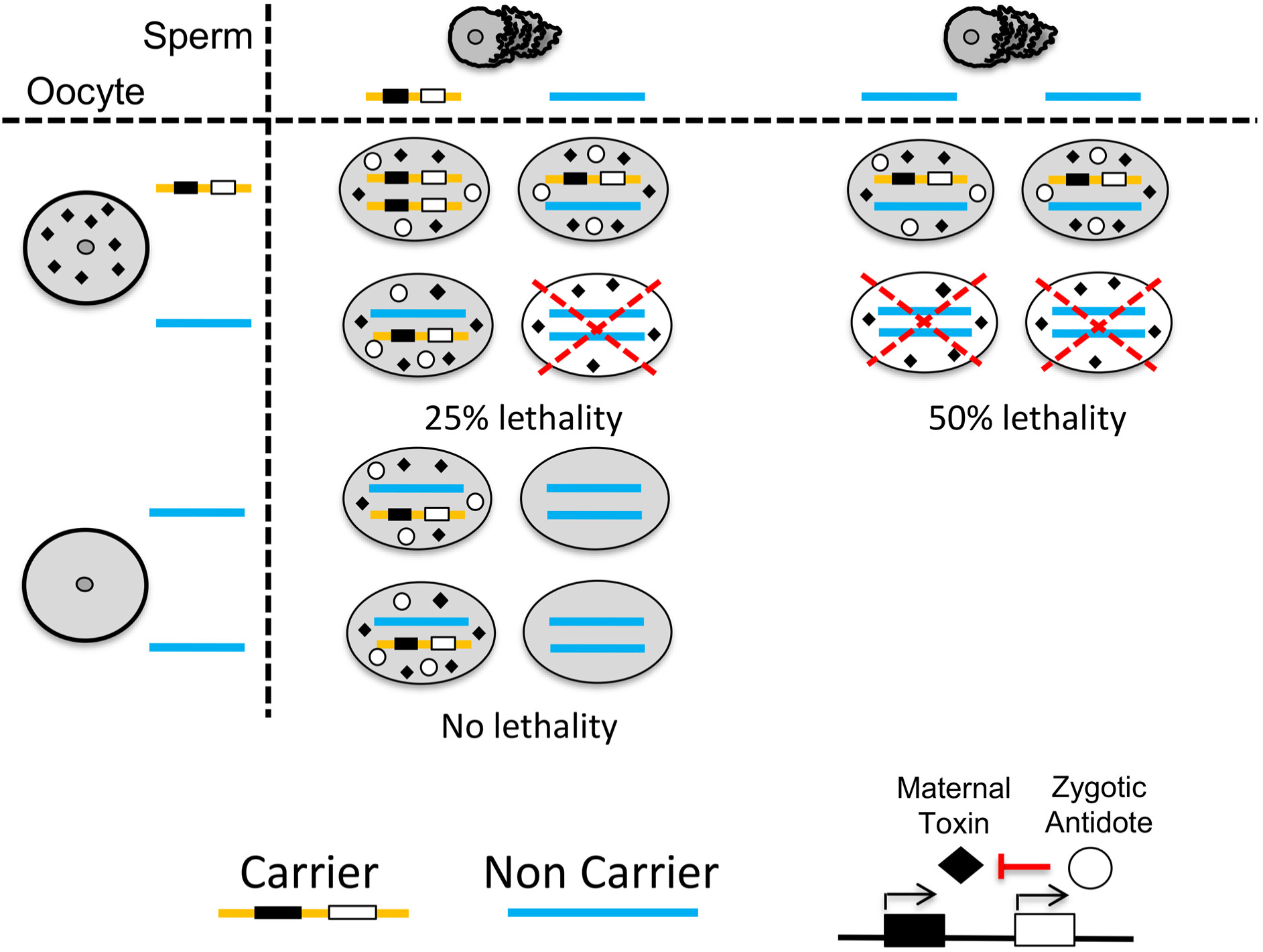
Expected F_2_ embryonic lethality due to the action of a maternal-effect toxin linked to its zygotic antidote. The selfish element (shown on the bottom right) is encoded by two tightly-linked genes: a maternal-effect toxin (black rhombus) and a zygotic antidote (white circle). In a cross between a carrier (orange) and a non-carrier (blue), all F_1_ will carry one copy of the element. When these F_1_ self (top left), all of their progeny inherit the maternal-effect toxin, regardless of their genotype. However, 25% of the progeny will not inherit a copy of the antidote, leading to lethality. A maternal-effect toxin can be distinguished from a paternal-effect toxin by backcrossing the F_1_ to non-carriers. When crossing F_1_ hermaphrodites to non-carrier males (top right), all of the progeny inherit the maternal-effect toxin, but 50% do not inherit the antidote and die. When the reciprocal backcross is done, none of the progeny inherit the toxin by maternal-effect, so all progeny that carry the toxin also carry the antidote, and no lethality is observed.

**Figure S3.**
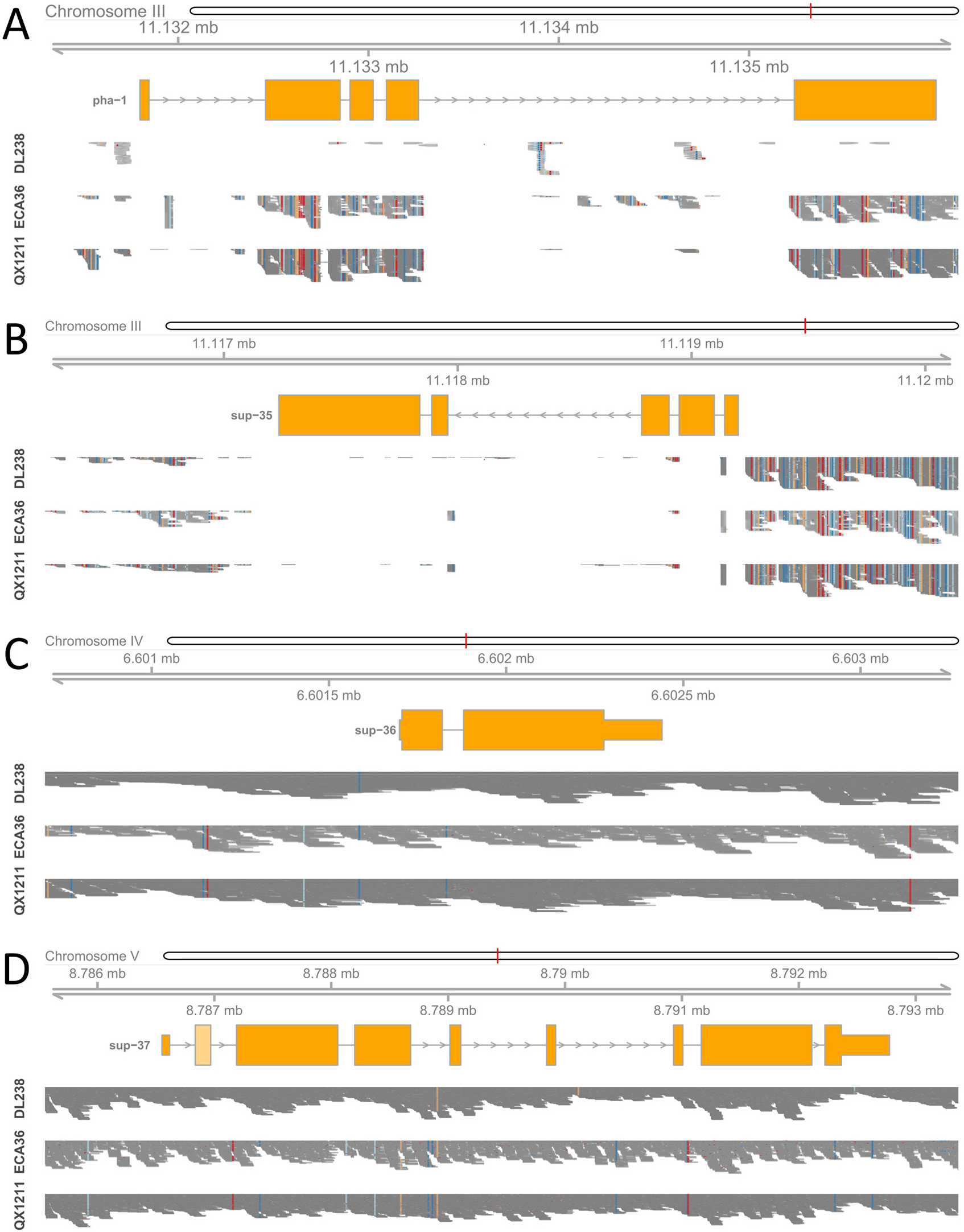
Alignment of short reads from DL238, ECA36 and QX1211 to the N2 reference in *pha-1, sup-35, sup-*36, and *sup-37*. Illumina short reads from DL238, ECA36, and QX1211 wild isolates were aligned against the N2 reference genome. A pileup of the reads aligning to *pha-1* **(A)**. *sup-35* **(B)**, *sup-36* **(C)** and *sup-37* **(D)** is shown. Gray bars illustrate a match and colors indicate mismatches. Lack of alignment strongly suggests that a gene is missing or that the sequence is highly divergent compared to N2.

**Figure S4.**
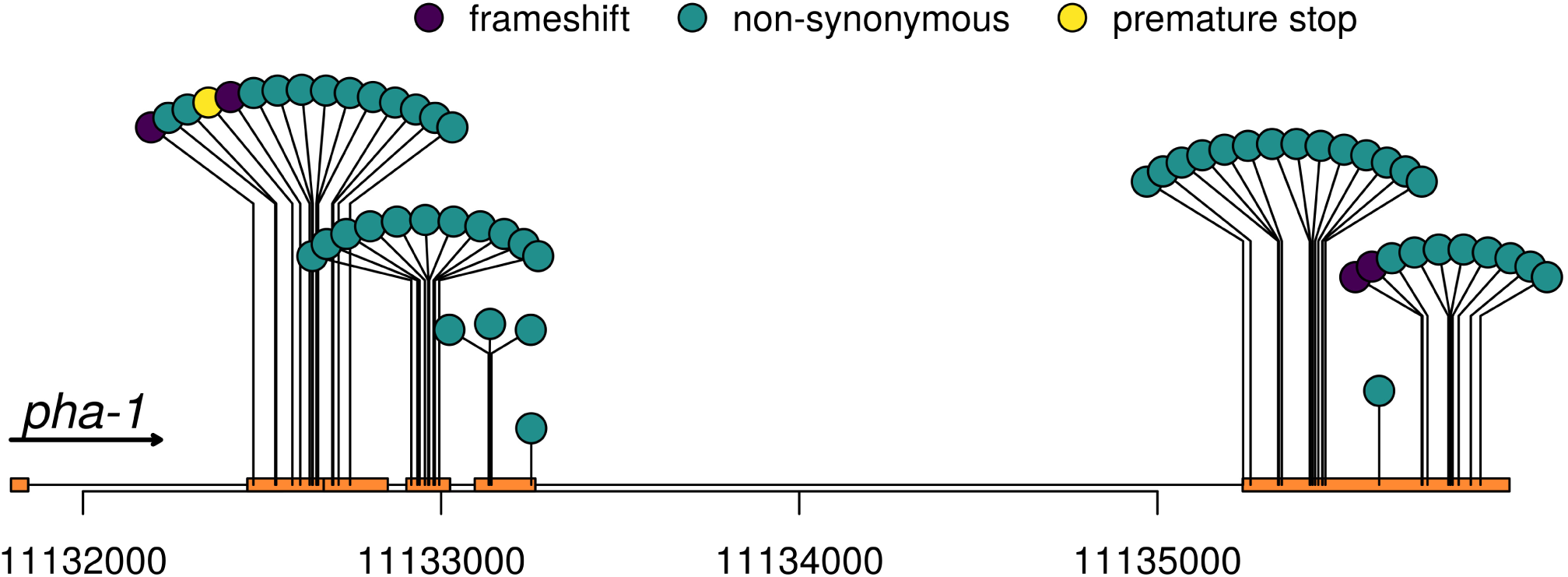
Variants in *pha*-1 in the QX1211 and ECA36 wild isolates. *pha-1* appears present in QX1211 and ECA36 but has accumulated a large number of mutations compared to the reference N2. The large accumulation of coding variants, including frameshifts, strongly suggests that *pha-1* underwent pseudogenization in QX1211 and ECA36.

**Figure S5.**
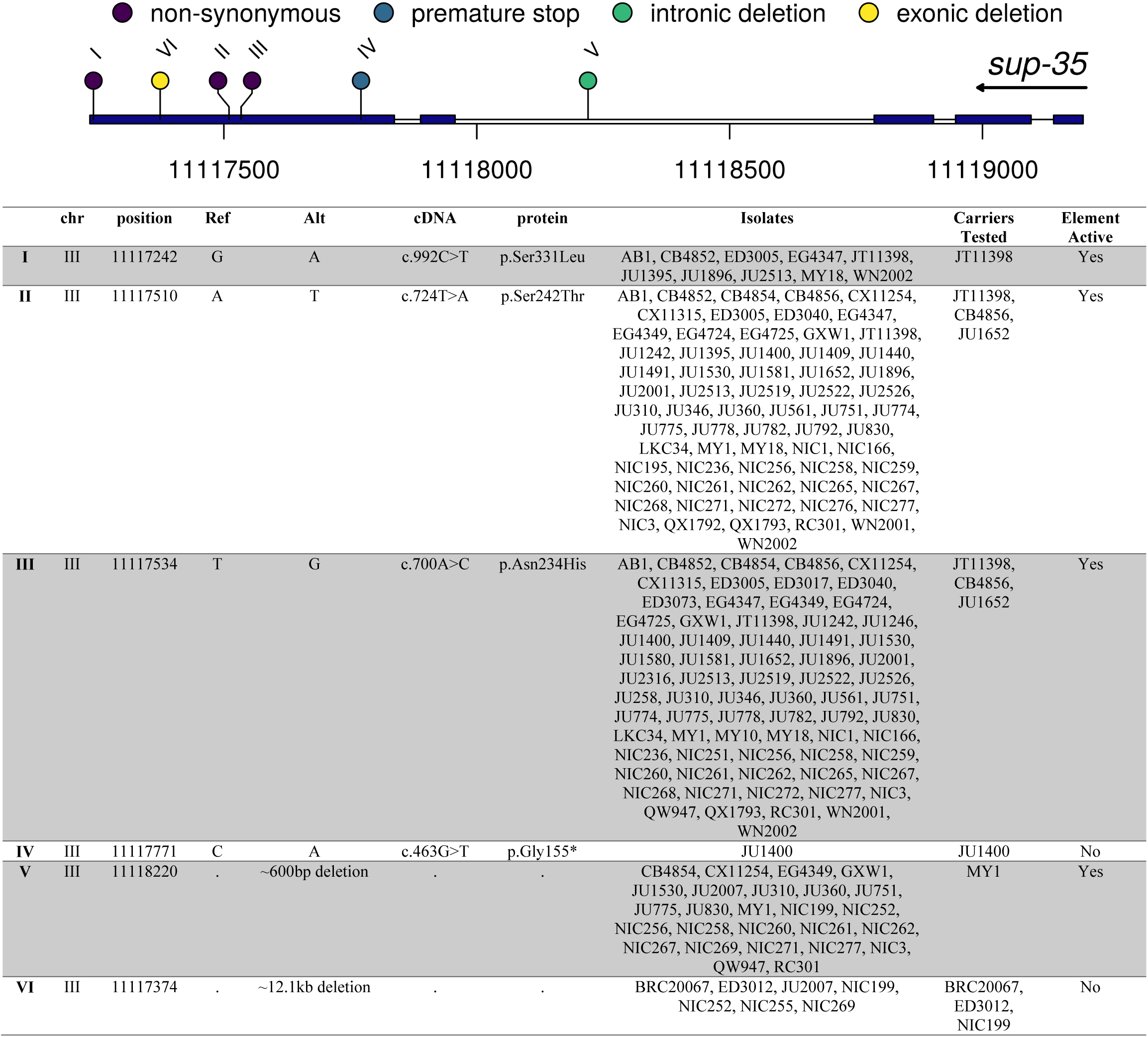
Coding and structural variants in *sup-35* across 152 C. *elegans* wild isolates. DL238, QX1211, and ECA36, which carry a pseudogenized copy of *sup-35* were excluded from the analysis. For each of the variants, we tested carriers by crossing them with DL238. In carriers of two variants (IV, VI), the maternal-effect incompatibility was abolished in the cross, indicating they are null *sup-35* alleles. For the intronic deletion (V), the center of the deletion is marked. For the large 12.kb deletion that includes part of the last exon of *sup-35,* the marker shows the position from which point *sup35* is deleted.

**Figure S6.**
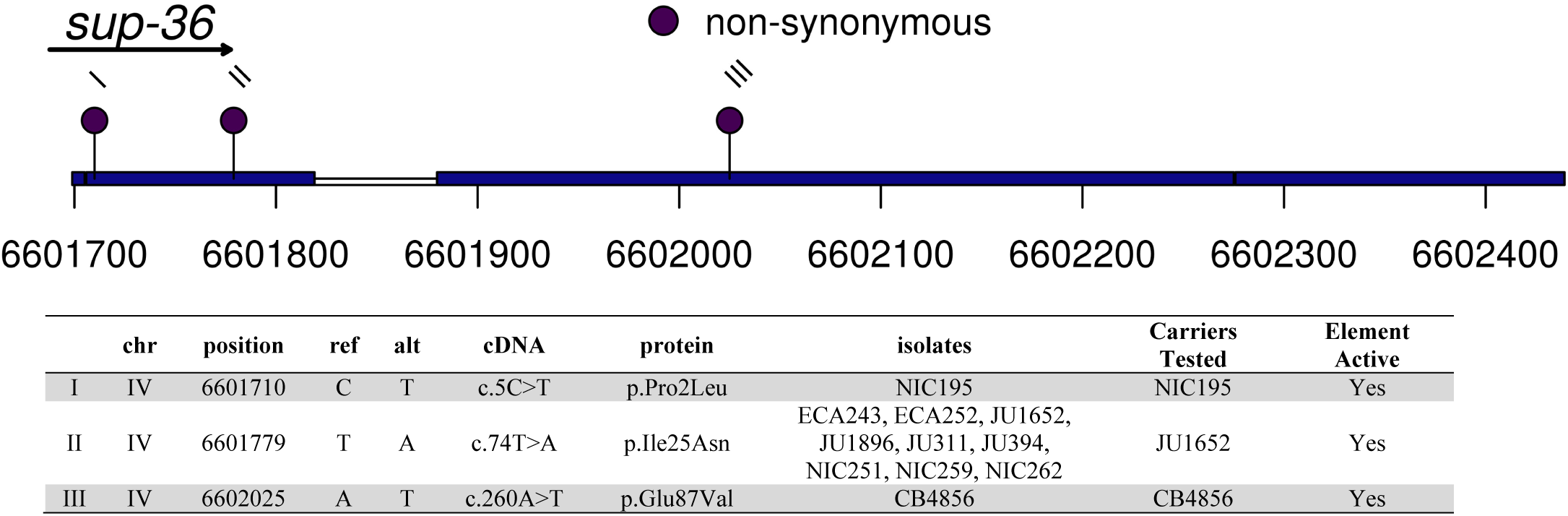
Coding variants in *sup-36* across 152 wild C. *elegans* wild isolates. Isolates carrying each variant were tested by crossing to DL238, and no evidence was found for reduced lethality.

**Figure S7.**
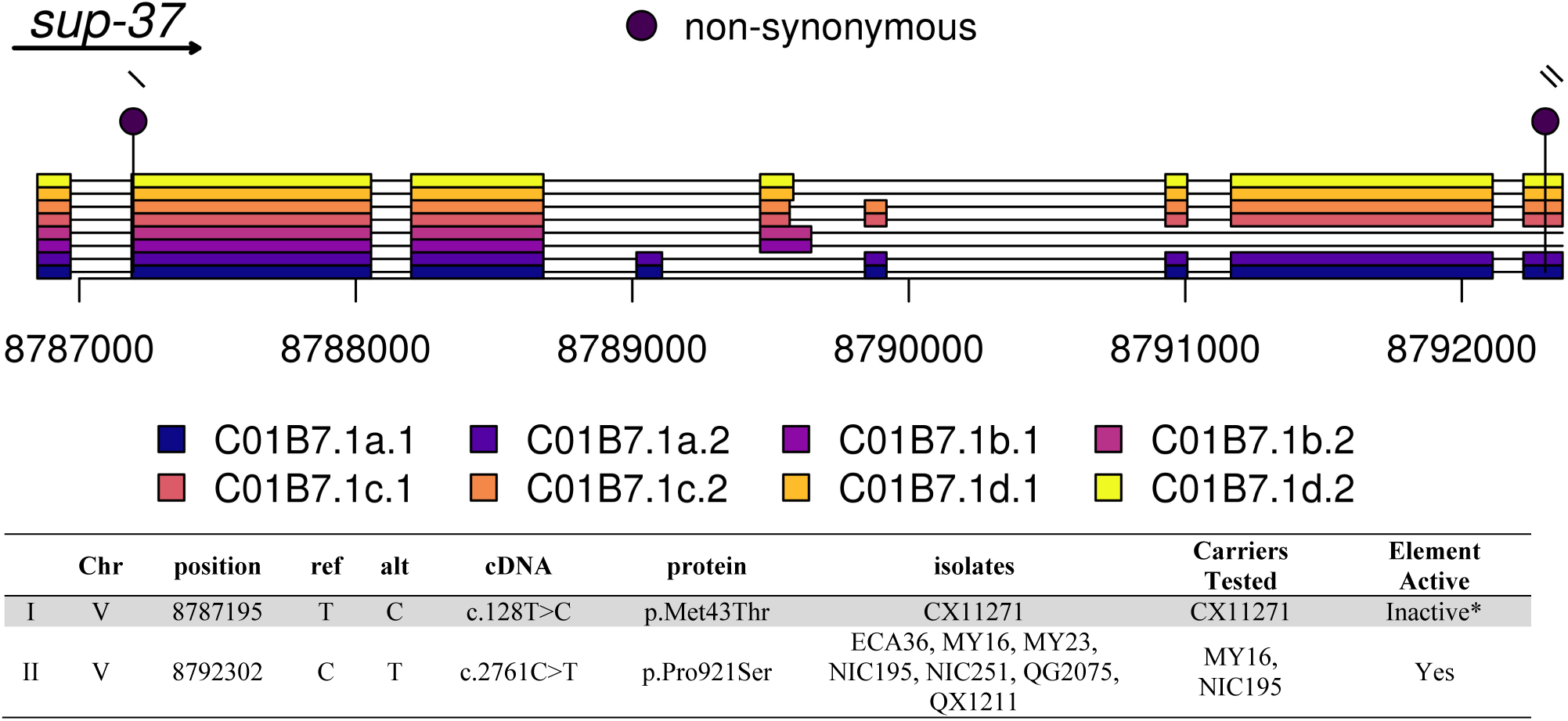
Coding variants in *sup-37* across 152 wild C. *elegans* wild isolates. Isolates carrying each variant were tested by crossing to DL238.

**Figure S8.**
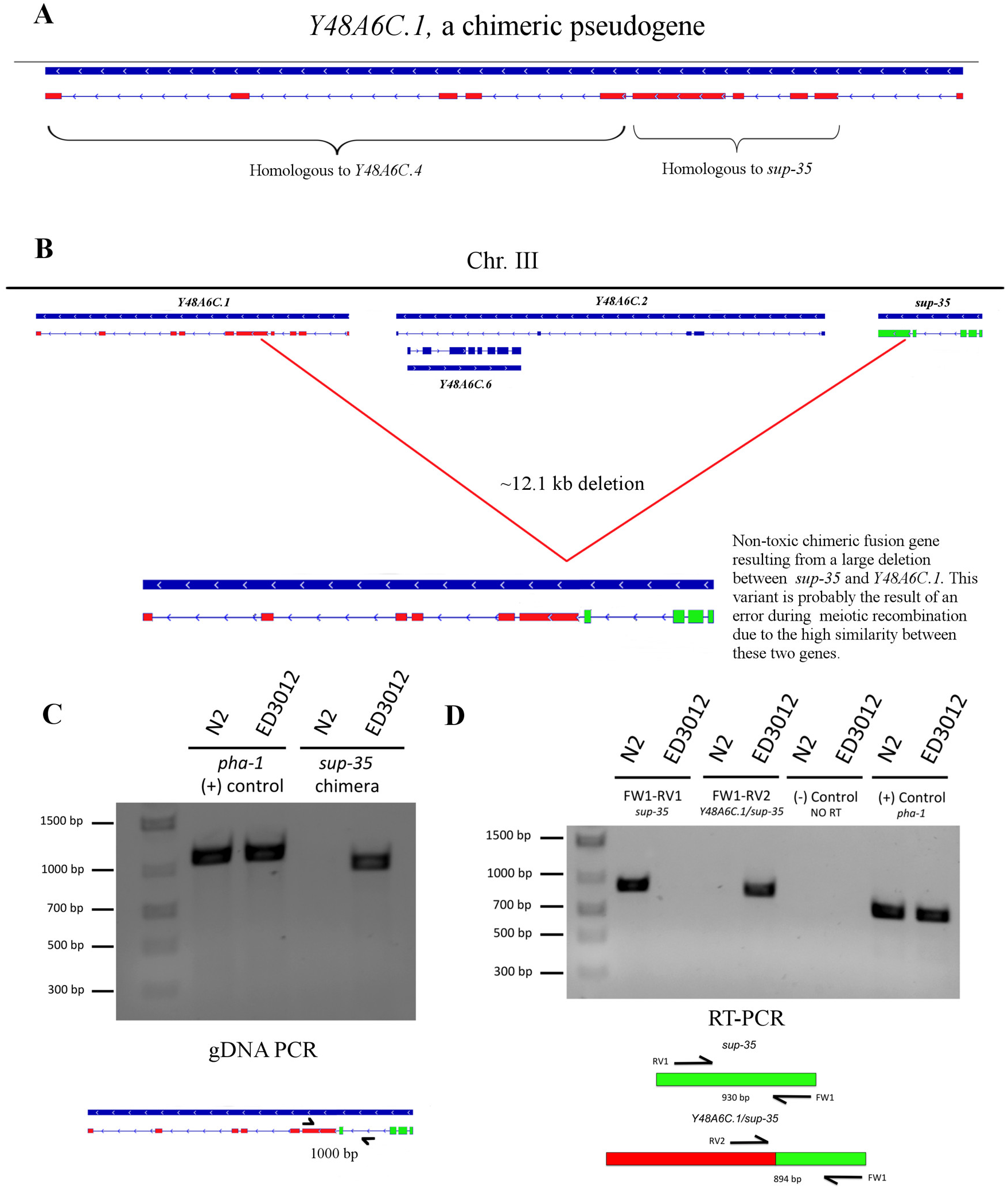
A large deletion identified in seven wild isolates results in a non-functional chimeric transcript that fuses *sup-35* and *Y48A6C.1.* *Y48A6C.1* is a pseudogene, which has four exons that are homologous to *sup-35* and five that are homologous to *Y48A6C.4.* **(B)** We identified 7 isolates (Fig. S4) that carry a 12.1kb deletion that fuses *sup-35* to *Y48A6C.1.* **(C)** Primers flanking the 12.1 Kb deletion amplify the expected amplicon in ED3012, a wild isolate carrying the deletion, but not in N2. **(D)** Reverse-transcription PCR (RT-PCR) from N2 and ED3012 worms shows that the chimeric *Y48A6C.1/sup-35* fusion gene is expressed and spliced. FW1/RV1 primer pairs are specific for the WT *sup-35* and FW1/RV2 are specific for the *sup-35/Y48A6C.1* fusion (FW1-RV2). Primer FW1 hybridizes in an exon-exon junction in *sup-35.* The RT enzyme was not added to the reaction as a negative control. *pha-1* expression was used as a positive control.

**Figure S9.**
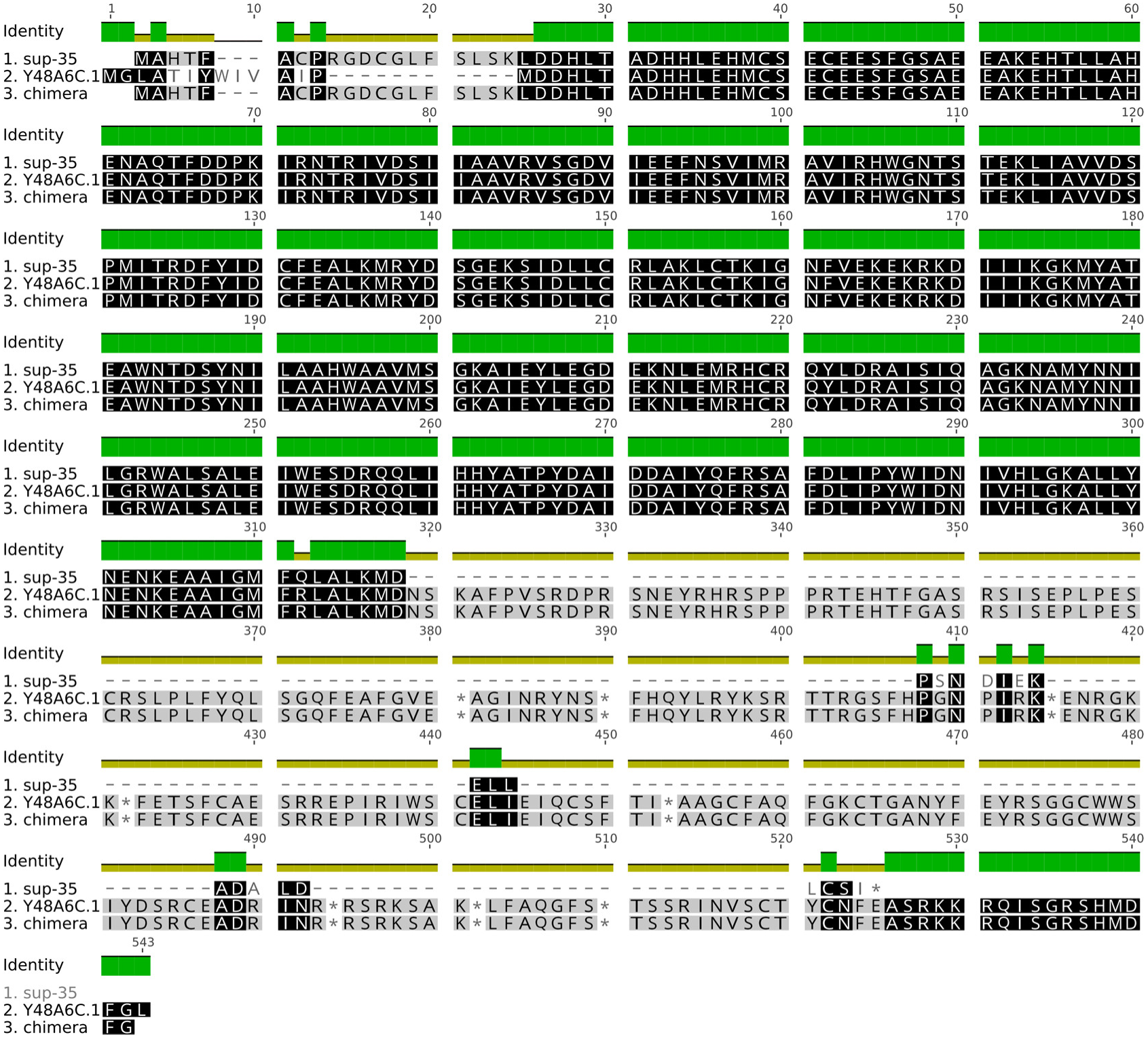
Protein alignment shows the homology between *sup-35,* the *Y48A6C.1,* and the chimeric gene that results from a deletion identified in seven wild isolates. Darker colors denote higher conservation. Similarly, conservation is also indicated by the color identity bar on top (red: low conservation, green: complete conservation. yellow: moderate conservation).

**Figure S10.**
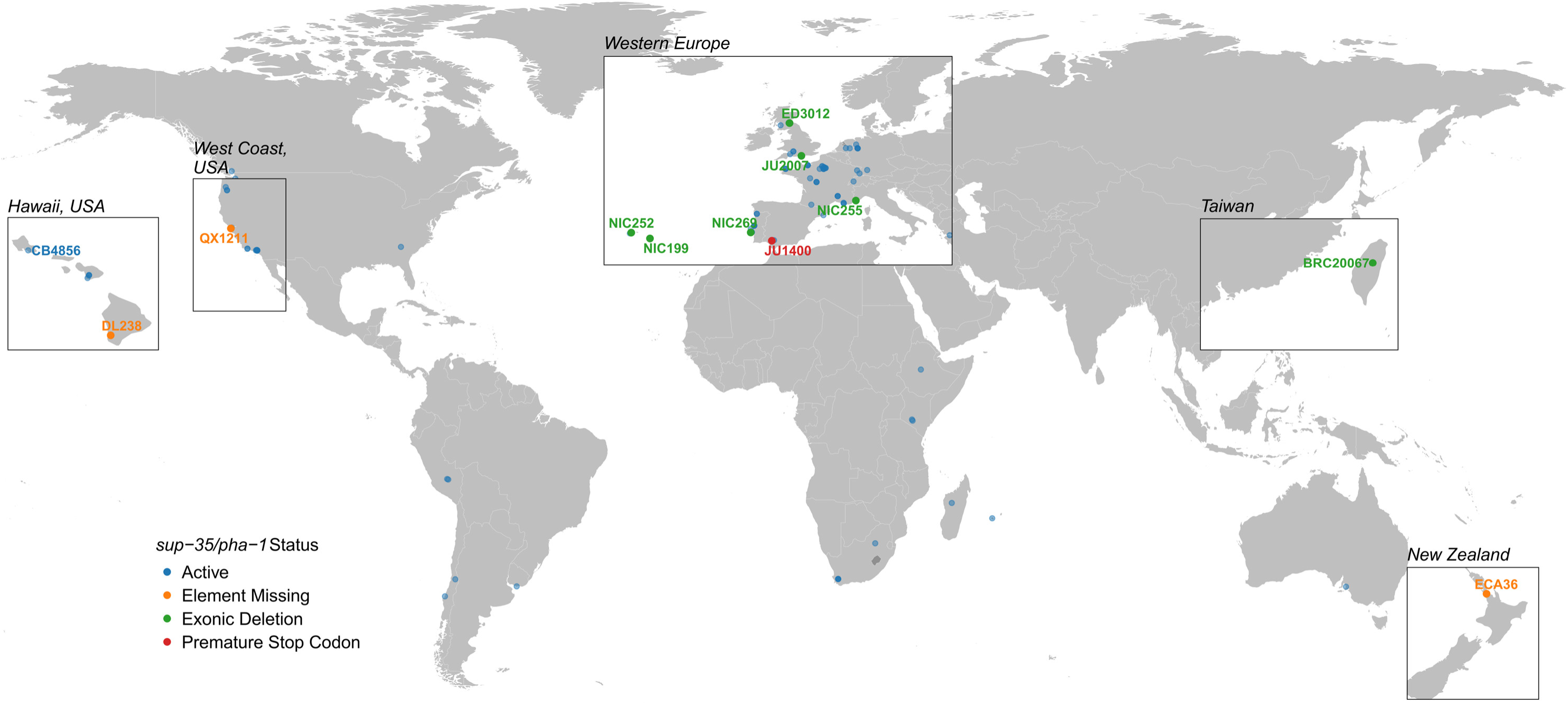
Worldwide distribution of *sup-35/pha-l* activity across 152 isolates. Each circle represents a single isolate. Only those isolates with an inactive *sup-35/pha-l* element are named, with the exception of CB4856, a highly divergent isolate from Hawaii, which nevertheless carries the active element.

**Figure S11.**
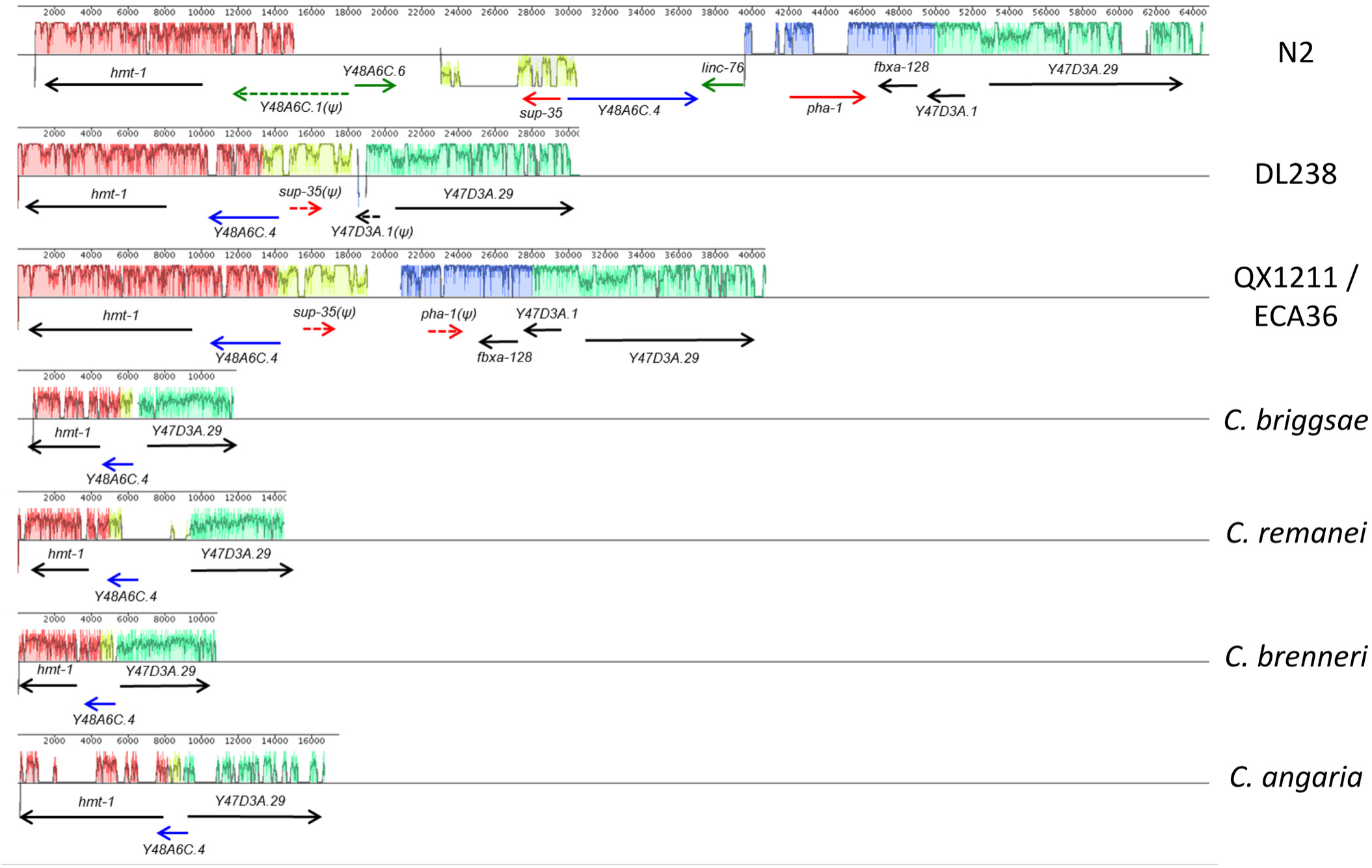
Multiple sequence alignment of the region containing the *sup-35/pha-1* element across *C. elegans* isolates and *Caenorhabditis* species. Two genes, *hmt-1* and *Y47D3A.29,* flank the *sup-35/pha-l element* in the N2 reference genome. Because these two genes are highly conserved in the *Caenorhabditis* genus, we performed a multiple sequence alignment of the genomic region located between these two genes using sequence derived from our de novo DL238 and QX1211 assemblies, as well as available genomes from other *Caenorhabditis* species. QX1211 and ECA36 share the same haplotype in this region. Both DL238 and QX1211 carry a pseudogenized version of *sup-35. pha-1* is missing from DL238 but pseudogenized in QX1211. In the four *Caenorhabditis* genomes available *(C. briggsae, C. remanei, C. brenneri* and *C. angaria),* the synteny of *hmt-1, Y48A6C.4,* and *Y47D3A.29* was conserved. The synteny was also conserved in DL238 and QX1211, but altered in the N2 haplotype. Based on this alignment, the most likely scenario is that the *sup-35/pha-1* element formed in the lineage leading to C. elegans. *sup-35* and *pha-1* were originally located next to each other in this ancestral haplotype and likely active. DL238, QX1211 and ECA36 carry a pseudogenized version of this ancestral haplotype. The ancestral haplotype also experienced an inversion affecting *sup-35* and *Y48A6C.4,* placing *Y48A6C.4* in between *sup-35* and *pha-1* and leading to the haplotype shared by the other 149 isolates tested, including N2.

**Figure S12.**
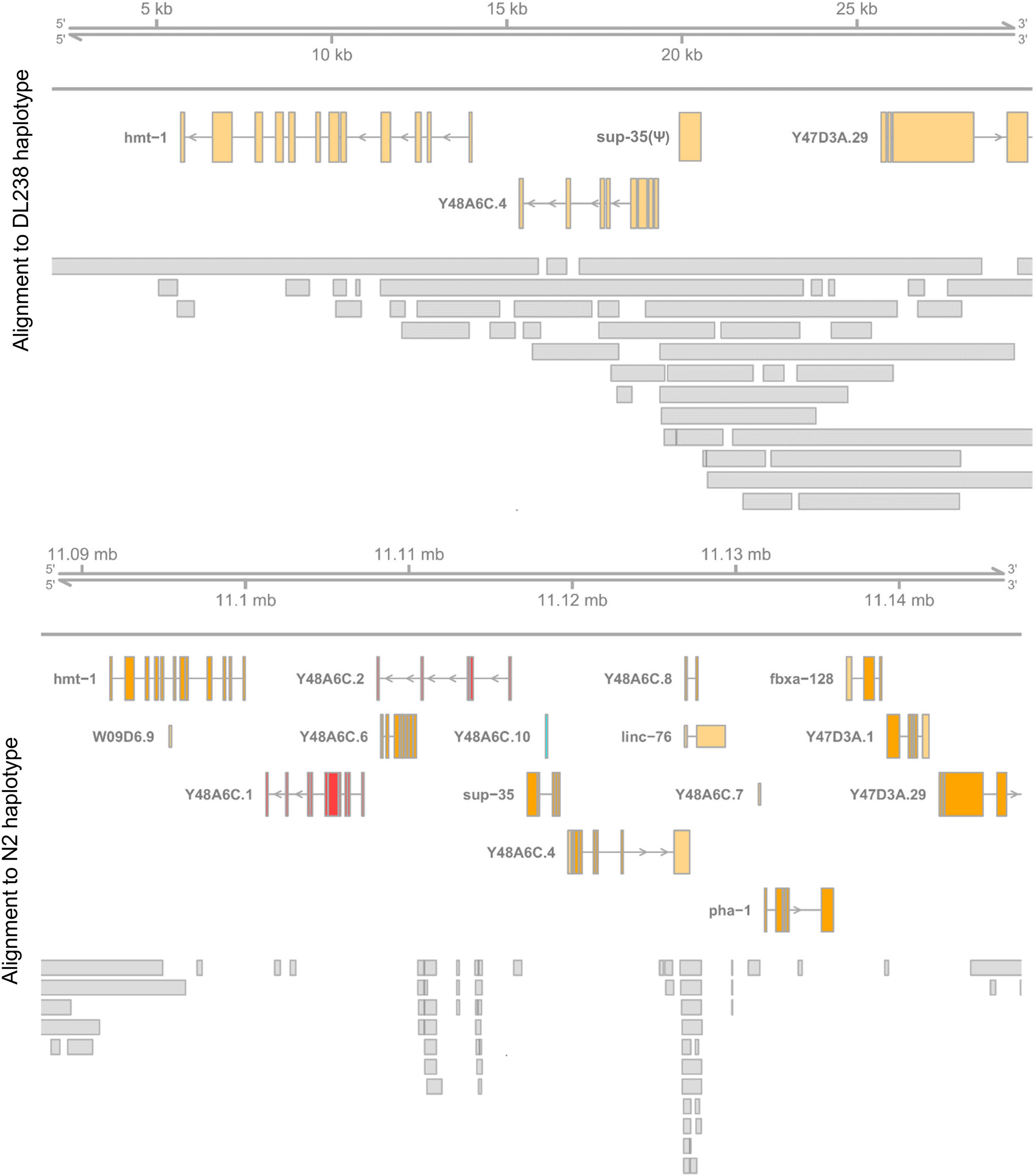
Alignment of DL238 Nanopore long read sequencing to the *de novo* assembled DL238 haplotype (top) and the N2 haplotype (bottom). Alignment to either the DL238 or the N2 haplotype is shown, along with the gene model in the region. *sup-35(Ψ)* is the pseudogenized version of *sup-35* found in DL238. The assembled DL238 haplotype is supported and independently validated by long reads that align across the region.

**Figure S13.**
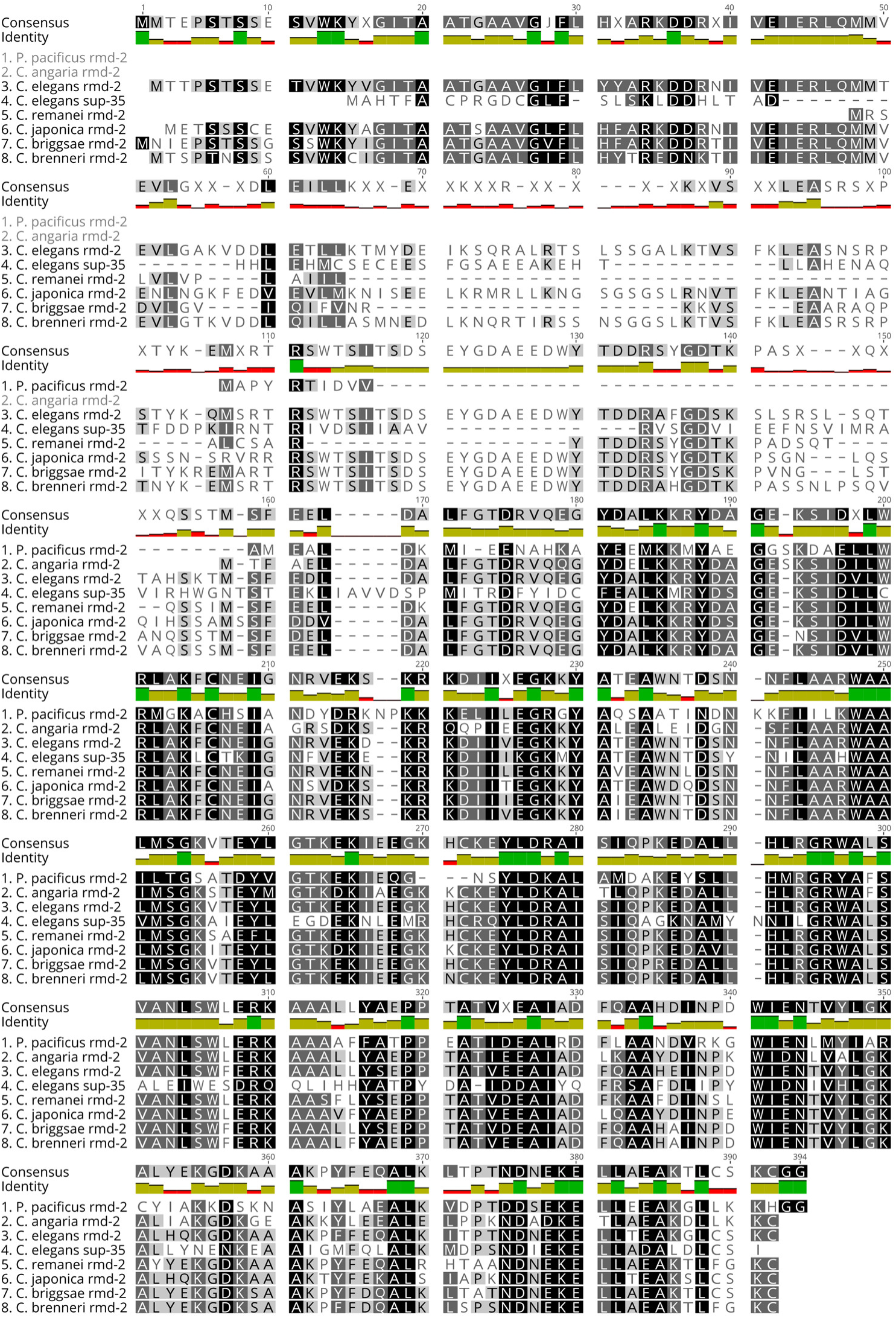
Protein alignment of *sup-35* against *rmd-2* from *Caenorhabditis.* Darker colors denote higher conservation. Similarly, conservation is also indicated by the color identity bar on top (red: low conservation, green: complete conservation. yellow: moderate conservation). For *C. elegans,* the largest transcript of *rmd-2 (C27H6.4b)* was used.

**Figure S14.**
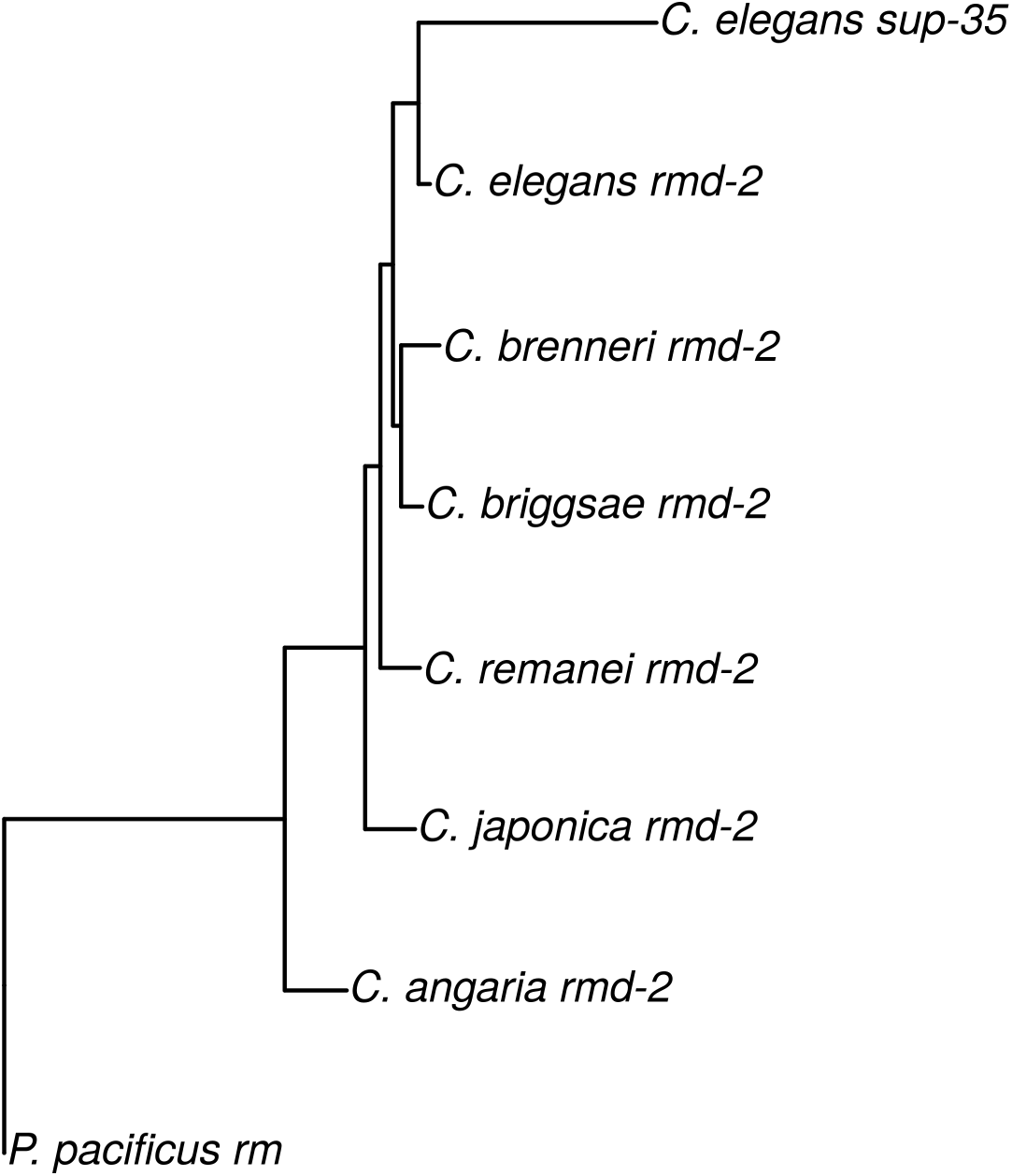
Phylogenetic tree of *rmd-2* and *sup-35* across diverse nematodes species. The phylogenetic tree for the cDNA of *rmd-2* was constructed as described for *Y48A6C.4. C.* elegans *rmd-2* is more closely related related to *sup-35* than to orthologous *rmd-2*, suggesting that *sup-35* arose from a duplication of *rmd-2* in the lineage leading to *C. elegans.*

**Table S1.** Screening for compatible isolates with DL238. Males from wild isolates carrying potentially functional variation in *sup-35, sup-36* or *sup-37* were crossed into DL238. The F_1_ progeny were allowed left to self, or were backcrossed into DL238. We backcrossed the F_1_ into either males or hermaphrodites to separate maternal-effect lethality from *sup-35/pha-1* from paternal-effect lethality from the *peel-1/zeel-1* element. The status of the *peel-1/zeel-1* element was also predicted by observing the sequencing in the isolates.

| Isolate | *peel-1/zeel-1*<br>Status | Seeded | Dead | Lethality | Seeded | Dead | Lethality | Seeded | Dead | Lethality |
| --- | --- | --- | --- | --- | --- | --- | --- | --- | --- | --- |
| **BRC20067** | Inactive | 450 | 0 | 0% | 210 | 10 | 4.76% | 164 | 1 | 0.61% |
| **ED3012** | Active | 417 | 97 | 23.26% | 245 | 115 | 46.94% | 276 | 12 | 4.35% |
| **JT11398** | Active | 215 | 88 | 40.93% | 175 | 95 | 54.29% | 305 | 141 | 46.23% |
| **JU1400** | Active | 733 | 176 | 24.01% | 123 | 68 | 55.28% | 185 | 5 | 2.70% |
| **JU1652** | Active | 110 | 41 | 37.27% | 185 | 90 | 48.65% | 184 | 88 | 47.83% |
| **MY1** | Inactive | 420 | 113 | 26.90% | 248 | 7 | 2.82% | 125 | 57 | 45.60% |
| **MY16** | Active | 499 | 198 | 39.68% | 182 | 100 | 54.95% | 404 | 169 | 41.83% |
| **NIC195** | Active | 377 | 170 | 45.09% | 232 | 127 | 54.74% | 166 | 73 | 43.98% |
| **NIC199** | Active | 207 | 55 | 26.57% | 153 | 79 | 51.63% | 183 | 5 | 2.73% |
| **CX11271** | Inactive | 689 | 117 | 17.00% | 238 | 1 | 0.4% | 490 | 197 | 42.82% |

**Table S2.**
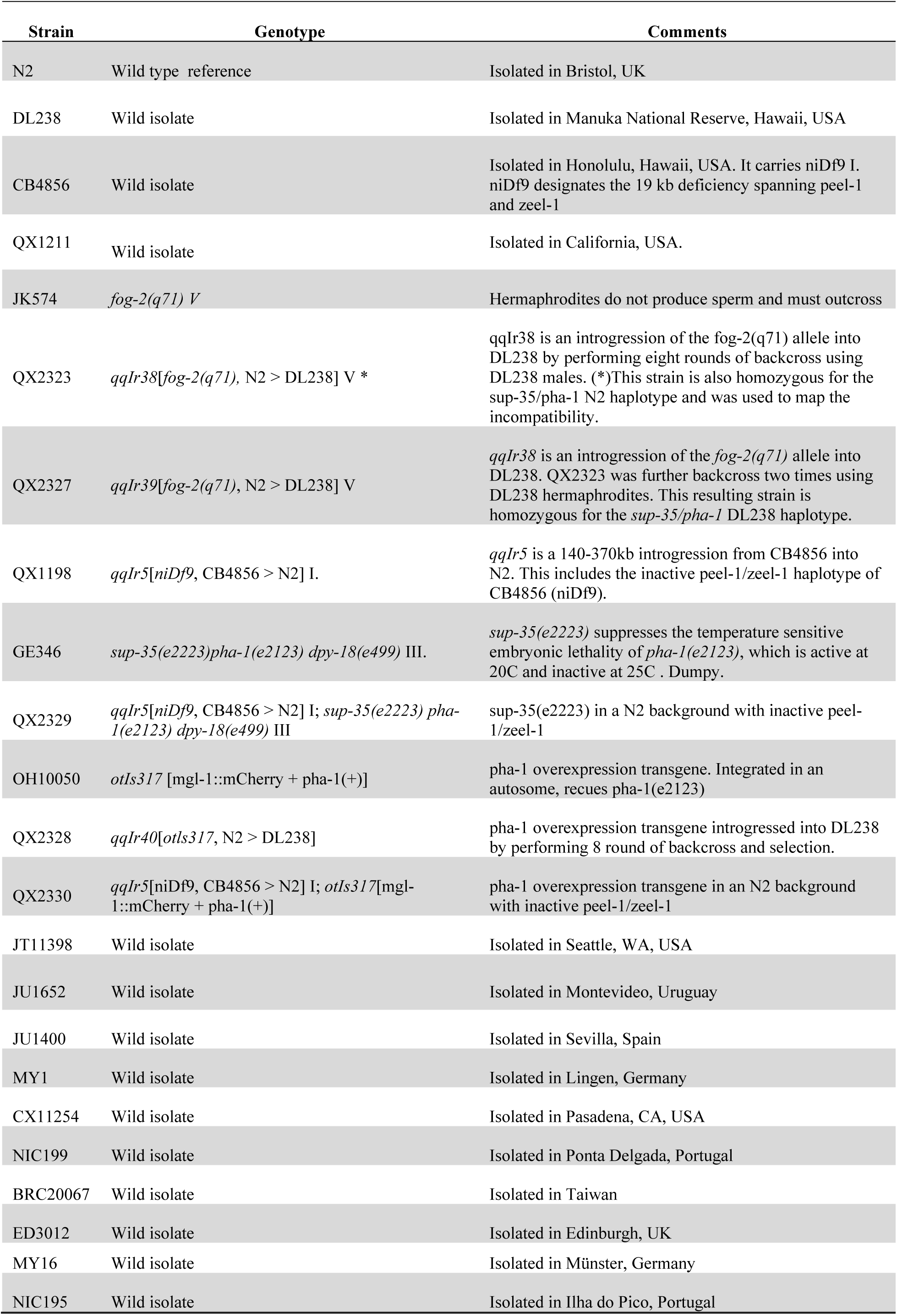
Strains used in the study.

